# ATM phosphorylates PP2A subunit A resulting in nuclear export and spatiotemporal regulation of the DNA damage response

**DOI:** 10.1101/2021.08.29.458108

**Authors:** Amrita Sule, Sarah E. Golding, Syed F. Farhan, James Watson, Mostafa H. Ahmed, Glen E. Kellogg, Tytus Bernas, Sean Koebley, Jason C. Reed, Lawrence F. Povirk, Kristoffer Valerie

## Abstract

Ataxia telangiectasia mutated (ATM) is a serine-threonine protein kinase and important regulator of the DNA damage response (DDR). One critical ATM target is the structural subunit A (PR65) of protein phosphatase 2A (PP2A), known to regulate diverse cellular processes such as mitosis and cell growth as well as dephosphorylating many proteins during the recovery from the DDR. We generated mouse embryonic fibroblasts expressing PR65-WT, -S401A (cannot be phosphorylated), and -S401D (phosphomimetic) transgenes. Significantly, S401 mutants exhibited extensive chromosomal aberrations, impaired DNA double-strand break (DSB) repair and underwent increased mitotic catastrophe after radiation. Our study demonstrates that the phosphorylation of a single, critical PR65 amino acid (S401) by ATM fundamentally controls the DDR, and balances DSB repair quality, cell survival and growth by spatiotemporal PR65 nuclear-cytoplasmic shuttling mediated by the nuclear export receptor CRM1.

## Introduction

Protein phosphatase 2A (PP2A) is a major serine/threonine (S/T) phosphatase that regulates many cellular processes including DNA replication, cell cycle progression, cell differentiation, and the DNA damage response (DDR) (1–3). The mammalian PP2A holoenzyme is a hetero-trimer (ABC) comprised of a scaffolding subunit (A or PR65), a regulatory subunit (B), and a catalytic subunit (C), or a heterodimer (AC; PP2A Core enzyme) (4). Both the A and C subunits have each two isoforms that are evolutionary conserved (5, 6). The two PR65 isoforms, Aα (PPP2R1A) and Aβ (PPP2R1B) are 86% identical. Each PR65 isoform is shaped into a horseshoe composed of 15 non-identical internal repeats of a 39 amino acid sequence, termed HEAT (<u>H</u>untingtin, <u>E</u>longation factor 3, Protein phosphatase 2A-subunit <u>A</u>, and the yeast kinase <u>T</u>OR1) motif (7–9). Almost 90% of the PP2A holoenzymes comprise of the Aα isoform whereas Aβ is primarily associated with development and the CNS (10). There are at least 17 subtypes of the B subunit, classified into four distinct families, B/PR55, B’/PR61, B’’/PR72 and PTP/PR53 (4,6,11,12). Specific regulatory B subunits control the diversity and target specificity of the various PP2A holoenzymes. The association of the AC heterodimer with a specific B subunit imparts complete activity, sub-cellular localization, and substrate specificity to the heterotrimeric holoenzyme.

Ataxia telangiectasia mutated (ATM), an S/T kinase, is defective in the human autosomal recessive disorder ataxia-telangiectasia (A-T) and is one of the major regulators of the cellular response to DNA damage (13). ATM is activated by DNA double-strand breaks (DSBs) induced endogenously by growth factor signaling, and exogenously by DNA damaging agents like ionizing radiation (IR) and radiomimetic drugs. In the event of DNA damage, ATM and other PIKKs such as A-T and RAD3-related (ATR), and DNA-PKcs get activated and direct a coordinated response referred to as the DDR by acting at several levels, including cell cycle checkpoints, DNA repair, and apoptosis (13–18). In a sense, ATM acts as a cellular rheostat balancing and coordinating the DDR with cell growth to maintain homeostasis (19). DDR proteins, phosphorylated in response to DNA damage are subsequently dephosphorylated by phosphatases to reverse the DDR and reset the cell to normalcy (14, 20). We have previously reported that the inhibition of ATM kinase blocks pro-survival signaling in human tumor cell lines by reducing AKT (S473) phosphorylation (21). This study revealed that ATM regulates AKT phosphorylation via an okadaic acid (OA) sensitive phosphatase. OA inhibits the S/T protein phosphatases protein phosphatase 1 (PP1) and PP2A, exhibiting greater affinity and inhibition towards the latter (22–24).

Protein phosphatase 2A regulates key DDR proteins ATM, ATR, CHK1, CHK2, and p53 (25–28), controls the G2/M checkpoint (29), and facilitates DSB repair via the dephosphorylation of γ-H2AX in response to IR (30). More specifically, PP2A binds to ATM and regulates ATM (S1981) auto-phosphorylation and activation in an indirect manner (25). Furthermore, it was reported that PP2A is regulated by ATM-mediated phosphorylation of PR65 at S401 resulting in the retention of histone deacetylase HDAC4 in the cytoplasm (31). Consequently, without functional ATM HDAC4 is unable to shuttle between the cytoplasm and nucleus where it permanently remains resulting in neurodegeneration which is a characteristic hallmark of A-T. Even though this study did not address the impact of PR65 S401 (de)phosphorylation during the DDR, a major conclusion from the report was that PP2A is negatively regulated by ATM, which does fit well with how we perceive ATM to regulate PP2A and AKT phosphorylation (21).

In the present study we generated PR65 depleted mouse embryonic fibroblasts (MEFs) expressing PR65 site-specific S401 mutants; a S401A mutant which cannot be phosphorylated as well as a phospho-mimetic S401D mutant. We found that these mutant cells were impaired in the DDR, displayed increased radio- and chemo-sensitivity, and had fundamentally altered DSB repair. Both S401 mutants were deficient in HRR with S401D also being severely compromised in non-homologous end joining (NHEJ) whereas S401A appeared to possess aberrant, greater than normal end joining activity. Upon assessing further, the phenotype of these cells, we uncovered a completely rewired DSB repair machinery due to PP2A malfunctioning. Here we report that the spatial and temporal regulation of PR65 phosphorylation at S401 by ATM through cytoplasmic-nuclear shuttling is critical for regulating DSB repair quality and the proliferative recovery from the DDR.

## MATERIAL AND METHODS

### Plasmids

Previously described pMIG-Aalpha WT retroviral vector with a Flag-tag at the NH_2_ terminal (32) obtained from Addgene (plasmid #10884) was generously provided by William Hahn. Mutations at the S401 codon and other positions were generated using QuikChange site-directed mutagenesis (Stratagene). The PR65 mutations were verified by DNA sequencing. Other plasmids used were Flag-hCRM1, a gift from Xin Wang (Addgene plasmid # 17647): pmCherry-C1-RanQ69L, a gift from Jay Brenman (Addgene plasmid # 30309): pCSCMV:tdTomato, a gift from Gerhart Ryffel (Addgene plasmid # 30530): human TERT (LOX-TERT-iresTK; Addgene plasmid #12245), generously provided by Didier Trono; DR-GFP plasmid (33); and pcDNA3-H2B-mCherry (Addgene plasmid #20972) (34, 35). PR65 was cloned into pcDNA3-mRuby2 (Addgene plasmid #40260), generously provided by Michael Lin, into pGEX-2T (GE Life Sciences #28954653) and pPAmCherry-a-tubulin (Addgene plasmid #31930) provided by Vladislav Verkhusha. Mutations at the S401 and L373 positions were generated using QuikChange site-directed mutagenesis (Stratagene). The PR65 mutations were verified by DNA sequencing.

### Antibodies and other reagents

Antibodies used for western blotting and/or immunocytochemistry were anti-ATM (Cat #2873), anti-AKT (Cat #2920), anti-p(S473) – AKT (Cat #9271), anti-PR65 (Cat #2039), anti-PP2A-C (Cat #2038), anti-pS139-(γ)-H2AX (Cat #9718) and anti-GST (Cat #2624) purchased from Cell Signaling Technology. Anti-γ-H2AX (Cat #05-636), anti-GAPDH (Cat #MAB374) were purchased from Millipore Sigma. Anti-53BP1 (Cat #NP100-304), anti-Rad51 (Cat #NB100-148), and anti-PyMT (Cat #NB100274955) were purchased from Novus Biologicals. Anti-p(S4/S8)-RPA (Bethyl laboratories, Cat #A300-245A), anti-ERK2 (Cat #sc-154), anti-pERK1/2 (Cat #sc-7383) and anti-CRM1 (Cat #sc-74454) were purchased from Santa Cruz Biotechnology, and anti-Flag (Sigma Aldrich, Cat #F1365). Alexa Fluor 594 goat-anti-mouse immunoglobulin G (IgG), Alexa Fluor 680 goat-anti-rabbit IgG, and Alexa Fluor 680 goat anti-mouse IgG (Life Technologies), Dylight 800 anti-mouse IgG (Cat #5257) and Dylight 800 anti-rabbit IgG (Cat #5151) were from Cell Signaling Technology. Leptomycin B (Cat #L2913) and camptothecin (Cat #7689-03-4) were from Sigma Aldrich. KU-60019 was from Selleck (Cat #S1570).

### Generation of PR65 conditional knockout mouse embryonic fibroblasts

All animal breeding and experiments were approved by Virginia Commonwealth University IACUC. Fvb.129S-*PPP2R1A^tm1wltr/j^* mice (36) were obtained from Jackson laboratories as stock number 017441. These CKO mice possess *lox*P sites flanking exons 5-6 of the PR65 gene (*PPPP2R1A*). Heterozygous 129S-*PPP2R1A^tm1wltr/j^* (CKO/WT) mice were bred to obtain 129S-*PPP2R1A^tm1wltr/tm1wltr^* (CKO/CKO) mice. Embryos were harvested at 13.5 dpc from a pregnant CKO/CKO female mouse and PR65-CKO MEFs generated by the VCU Massey Cancer Center Transgenic/Knockout Mouse Core using standard methods (37).

### Cell culture and irradiation

HEK293, U1242, U87 glioma cells, and MEFs were cultured in DMEM (GIBCO) medium supplemented with 10% FBS and 1% Pen-Strep (Life Technologies). Serum starvation was for 16 hours (wherever mentioned) in medium without FBS. Irradiations were done using an MDS Nordion Gammacell-40 research irradiator with 137-Cs source delivering a dose rate of 1.05 Gy/min. Cell growth was determined by CellTiter-Glo® Luminescent Cell Viability Assay (Promega, Cat #G7570). Cells were serially diluted and seeded in a 96-well plate. At days 2, 5 and 7 after seeding, CellTiter-Glo® reagent was added to the medium at the recommended final concentration. Plates were incubated for 10 minutes at room temperature and luminescence determined using an EnVision Multilabel reader (PerkinElmer) and the readings taken as the measure of cell growth. Clonogenic survival assays were carried out as described (38, 39).

### Microscopy and cell imaging

For DNA repair foci assay, cells were grown on Lab-Tek (Naperville, IL) glass slides. After treatment, cells were fixed with 3% paraformaldehyde, made permeable with 0.5% Triton-X-100 in phosphate buffered saline (PBS) and blocked with casein (ThermoFisher Scientific Cat #37528). Cells were stained with primary antibodies diluted in casein blocker followed by secondary antibodies, and nuclei were counterstained with DAPI (1 μg/ml) and mounted in Vectashield® mounting medium (Vector Labs). Cells were imaged and analyzed using Zeiss LSM 710 imaging system in the VCU Microscopy Facility.

For photoactivation studies to capture representative images, HEK293 cells were transfected with pPAmCherry-PR65 (WT, S401A and S401D) in glass bottom dishes and 48 h post-transfection sub-nuclear ROI (4 μm x 4 μm), marked by the squares, were photoactivated by 405 nm laser (242.8 μJ) and followed over time (<300 s) by live cell imaging using a Zeiss LSM 710 microscope with images acquired at 63x. For quantitative photoactivation studies, HEK293 cells, stably transfected with linearized pPAmCherry-PR65 (WT, S401A and S401D, NES, S401A-NES and S401D-NES), were cultured under G418 (500 μg/ml) selection in a glass bottom dish for 48 h. Next, the cells were imaged with Zeiss LSM 880 confocal system, based on Axio Z1 inverted stage and equipped with 40x PlanApo oil immersion objective (NA = 1.4), 405 nm diode laser (30 mW) and 561 nm DPSS laser (20 mW). Fluorescence of mCherry was excited with the latter laser attenuated to 0.8% and detected in 570 – 620 nm range with a multi-anode (spectral) GaAsP hybrid detector in the integrative mode at 950V gain. Time series of transmitted light and fluorescence images were collected with 0.28 µm pixel size, 3.07 µs dwell time (236 µs frame time) and 256x256 pixel frame size. Following first 25 frames 2.8x2.8 µm regions in cell nuclei were irradiated with 325.2 µJ of 405 nm light. Next, imaging mCherry fluorescence resumed for 277 s (1175 frames). Up to 7 irradiation regions were selected in each field of view using transmitted light image as the reference.

### *In vitro* kinase assay

YFP-ATM was immunoprecipitated from stably transfected HEK293 cells by GFP-TRAP (Chromotek). The immunoprecipitates were suspended in kinase assay buffer containing 2 μCi of [γ-^32^P] ATP, 20 μM unlabeled ATP, and GST-substrates (GST-PR65 or GST-p53). GST-PR65 has GST fused to full-length PR65 and GST-p53 expresses the first 100 amino acids of p53 (40). GST-fusions were purified from extracts of BL21 (DE3)/pLysS cells using Glutathione Sepharose 4B beads as recommended by the manufacturer (GE Healthcare Life Sciences). The kinase reaction was conducted at 30°C for 20 minutes and stopped by adding 2X Laemmli loading buffer. Samples were separated on 10% polyacrylamide gels, followed by drying of the gel onto 3M paper, exposed to a screen, scanned by a Phosphorimager (Typhoon) and the signal quantified (40).

### Transfection

Superfect, Effectene (Qiagen) or PEI were used to transfect cells transiently and for stable drug selection. Electroporation of MEFs was done using an Amaxa Nucleofector apparatus. Sterile Solution I: 20% ATP-disodium salt, 12% MgCl_2_ (Filter Sterilize, adjust pH to 7.4) and Solution II: 2% KH_2_PO_4_, 2% NaHCO_3_, 0.06% glucose (pH to 7.4) were combined 1:50 (total volume -100 μl) prior to electroporation (41). Electroporation was done with program: N-024.

### DNA repair assays

pEGFP-C1 (Clontech) plasmid was digested with BamHI and BglII to generate a 51-bp fragment which was inserted into the BamHI site of pCSCMV:tdTomato to generate a SmaI restriction site immediately downstream from the BamHI site. The modified pCSCMV:tdTomato + linker plasmid was linearized with SmaI followed by BamHI to create incompatible DNA ends and gel purified prior to use. pCSCMV:tdTomato+linker plasmid (1μg) was transfected into the MEFs (from a 80% confluent 6-cm plate) and cells divided equally into the wells of a 12-well plate. Cells were collected at 2, 4 and 8 hours from the media and pooled with the remaining cells after trypsinization. A small fraction was used to determine RFP+ cell numbers using a Nexcelom cellometer (Nexcelom Bioscience). Remaining cell pellets were re-suspended in 15% sucrose, 50 mM Tris-HCl, 25 mM EDTA buffer supplemented with 0.1% TritonX-100 and treated with RNase. Cell lysates were incubated at 50°C for 3-4 hours. Equal amount of re-suspension buffer was combined with equal volume of 0.1% SDS and 5 μg/ml of Proteinase K. Samples were incubated at 60°C overnight. This was followed by phenol chloroform treatment and DNA was precipitated with 1:1 isopropanol. The level of repaired DNA was assessed by quantitative real-time PCR, using an ABI 7900HT (Applied Biosystems) and SYBR green detection. The (Forward 5’ CGGAGCAAGCTTGATTTAGGTG 3’ and Reverse 5’ CGCATGAACTCTTTGATGACCTC 3’) primers were used to analyze the repair junction between BamHI and SmaI with primers for ampicillin gene (Forward 5’TGTGCAAAAAAGCGGTTAGCT 3’ and Reverse 5’GCGGCCAACTTACTTCTGACA 3’) were used for template normalization. MEFs were arrested in mitosis by treating cells with 50 ng/ml of nocodazole for 16 hours. Cells were collected for western blotting at 1, 2 and 4 hours after nocodazole washout with p-PLK1 (T210). MEFs were treated with 1 mM of hydroxyurea (HU) for 2 hours followed by a washout of the drug and cells collected at 0, 1, and 6 hours after washout and western blotting with antibodies specific for pRPA (S4/S8) and γ-H2AX.

In vitro end joining assay was carried out at 37°C in 50 mM triethanolammonium-acetate pH 7.5, 1 mM ATP, 1 mM dithiothreitol, 50 μg/ml BSA, 1.3 mM magnesium acetate and dNTPs at 100 μM each. Typically, a 16-µl reaction contained 10 µl of whole-cell extract (44), resulting in a final concentration of 8 mg/ml protein, 66 mM potassium acetate and 16% glycerol, and an effective Mg^2+^ concentration of 1 mM. The substrate was an internally labeled plasmid with partially cohesive ends, prepared as described (42, 43). Homologous recombination GFP assay (DR-GFP) was carried out as previously described by determining GFP+ cells by flow cytometry (44, 45). PR65^flox/flox^

MEFs were stably transfected with linearized DR-GFP plasmid and selection with puromycin followed by infection with WT, S401A or S401D pMIG-Aα (PR65) lentivirus. Pooled GFP+ cells were then infected with Ad-Cre to remove the endogenous PR65 (see **Fig. 1**).

**Figure 1.**
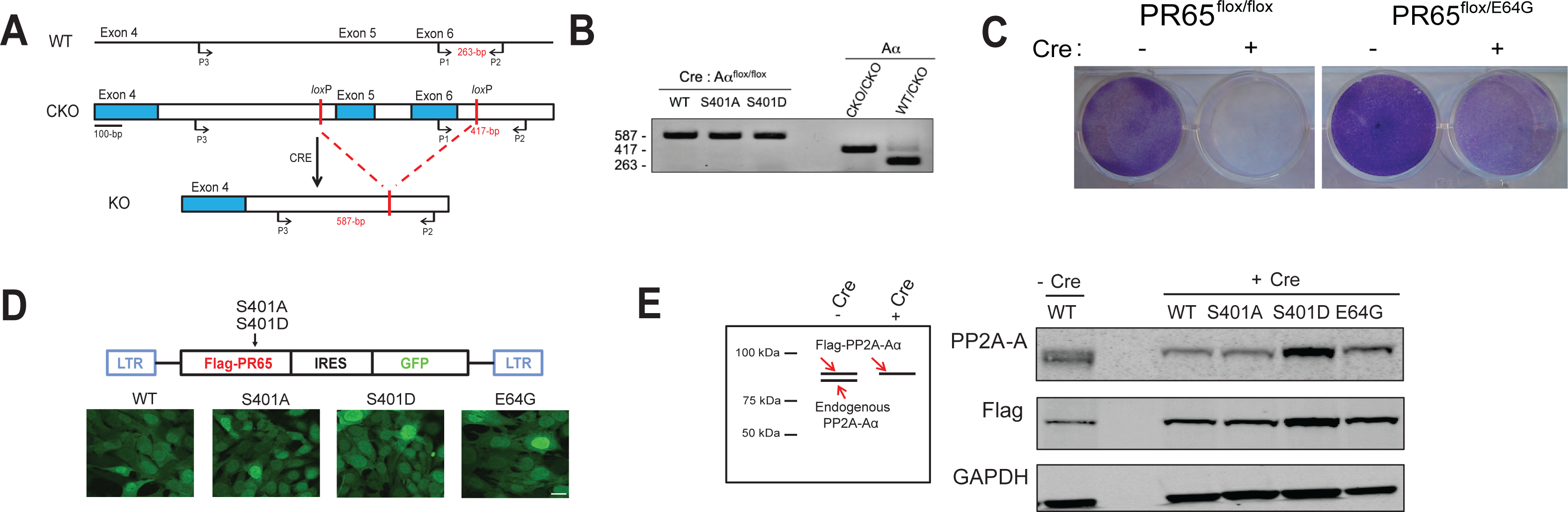
Generation of PR65 KO MEFs complemented with S401 mutants. **(A)** Targeting strategy for generation PR65 conditional knockout (CKO) MEFs. Expression of Cre recombinase in these PR65-CKO MEFs would generate a PR65 KO MEFs by deletion of exon 5 and 6. PR65-CKO MEFs were infected with retroviral vector pMIG-Aalpha, expressing Flag-tagged PR65 WT, S401A or S401D mutant in addition to GFP. This was followed by Infection of cells with Ad-CMV-Cre recombinase to knock out the endogenous PR65 alleles. **(B)** Genomic DNA was isolated from PR65 KO MEFs (WT, S401A and S401D) and PCR amplified. PCR screening with primers P1, P2, and P3 shown in **(A)** generated a 263-bp product for WT allele and a 417-bp product for the CKO allele. In cells where Cre was expressed, primer pair P2 and P3 generated a 587-bp product for the KO PR65 allele. **(C)** PR65 is essential for the survival of MEFs. PR65 CKO MEFs were infected with Ad-CMV-Cre recombinase or not. Cells were stained with crystal violet after one week to assess cell viability. PR65-CKO MEFs infected with Ad-CMV-Cre recombinase did not survive whereas PR65^flox/E64G^ MEFs did. **(D)** Plasmid map of pMIG-Aalpha WT with mutations marked (*top*) and expression of GFP in WT, S401A, S401D, and E64G MEFs with endogenous *PR65* alleles KO (*bottom*). Scale bar is 10 μm. **(E)** Western blot verifying the Flag-PR65 expression of site-specific mutants and floxing of endogenous PR65 in MEFs. **Left panel:** Explanation of the protein bands seen by western blotting. **Right panel:** Whole cell extracts of GFP+ sorted PR65 WT, S401A and S401D cells infected with Ad-Cre and uninfected PR65-WT MEFs were separated on an SDS-PAGE gel and analyzed by western blotting using anti-PR65 antibody, anti-Flag and anti-GAPDH (loading control) antibodies.

CRISPR-Cas9 DSB repair and chromosomal translocation assay was performed as described (46, 47). PR65-KO MEFs (WT, S401A and S401D) were infected with lentivirus expressing Cas9 and gRNA (Rosa26) and (Rosa26; H3f3b), respectively, by introducing one or two guide RNAs (5′-GTTGGCTCGCCGGATACGGG-3′ for H3f3b; 5′-ACTCCAGTCTTTCTAGAAGA-3′ for Rosa26) into LentiCRISPR v2 (Addgene plasmid #52961) and LentiCRISPRv2 hygro (Addgene Plasmid #98291). Plasmids were sequenced to confirm the correct gRNA insertions. To generate mixes of stable cell populations lentivirus was made in HEK293T cells and MEFs infected followed by selection with puromycin or hygromycin as described (48, 49). The Rosa26 and H3f3b primers used for PCR were those described (47). Amplified products were cloned in pGEM®-T Easy (Promega) and sequenced to verify translocation and junction sequences.

Chromosome translocation and amplicon length heterogeneity were quantified using high-speed atomic microscopy (HSAFM) with sample preparation and execution of the technique described in detail previously (50–52). Briefly, a tip with a <10 nm sharpened point was rastered across a surface, and the height of the tip measured to yield a three-dimensional map of the sample features with nanoscale resolution (53). Our high-speed atomic force microscope (Bristol Nanodynamics) routinely acquires 2 μm x 2 μm images with single-nanometer resolution at a rate of 1 image per second (54). To deposit DNA onto an atomically flat surface for imaging, we first purified the DNA from PCR reactions using Ampure (Beckman Coulter), then added MgCl_2_ to facilitate adhesion of DNA to negatively-charged mica. The resulting DNA solution was drop-deposited onto freshly cleaved mica (Ted Pella, Inc), dried with compressed air, and baked for 10 minutes at 120°C. Following HSAFM imaging, we employed a custom line-by-line processing program to flatten the images, which is required to account for mechanical and thermal vibration. To quantify DNA lengths observed in the flattened images, we used custom computer vision programming to automate DNA tracing and filter out strands that did not meet certain quality metrics, e.g. strands that were branched or circular. Analysis of traced length distributions was performed by fitting Gaussian mixture models, then analyzing the Bayesian information criteria to determine the number of Gaussian distributions that produced the best fit. The 95% confidence interval for the mean of each fitted Gaussian was determined using a bootstrapping approach with *n =* 10,000. All programming and analyses were conducted using MATLAB.

### Molecular modeling studies

Molecular Dynamics (MD) simulations were carried out with the NAMD 2.8 package developed by the theoretical and computational biophysics group in the Beckman institute for Advanced Science and Technology at the University of Illinois at Urbana-Champaign (55). CHARMM (Charmm-27) was used as the force field (56). The initial MD setup was done using VMD (57). Both S401 phosphorylated and unphosphorylated PR65 3D models were solvated in an equilibrated TIP3P water box. Then Cl^-^ and Na^+^ ions were added to neutralize the system and appropriate number of ions added up to a concentration of 50 mM. Solvent molecules were first minimized for 500 steps of conjugate gradients minimization method, keeping the protein molecules fixed to allow favorable distribution of water molecules on the complex surface. Subsequently, the system was coupled to a heat bath from 0 to 300°K and the constraints applied to the solute atoms were gradually decreased after which, the system was allowed to be simulated without restraints for over a period of 10 ps.

Finally, a 50 ns molecular dynamics production phase was carried out on the entire system. The analysis of the MD trajectory was done in VMD (56). HINT analysis was carried out on published x-ray crystal structures of the Holo and Apo PR65 protein (PDB IDs 2NPP and 1B3U respectively), which were also used for the docking studies with the CRM1 protein (PDB ID 3GB8). Protein-protein docking was performed using the HADDOCK (High Ambiguity Driven DOCKing) algorithm (58). The residues from CRM1 forming the protein-protein binding interface with Snurportin-1 were used as constraints for docking together with the identified putative leucine-rich nuclear export signal on PR65 (59, 60). Docking results were individually inspected after which high scoring models were passed into the refinement step. All docked poses were refined with the FireDock (Fast Interaction Refinement in a molecular Docking) algorithm (61), rescored and ranked using the HINT force field that describes and quantifies all interactions in the biological environment by exploiting the interaction information implicit in logP_o/w_ (the partition coefficient for 1-octanol/water solute transfer) (62, 63). HINT is known as a “natural” force field because it is based on empirical energetic terms that are defined by real experiments, and thus encodes interaction types including Coulombic, hydrogen bond and hydrophobic interactions expected between molecules in the biological environment as a free energy force field that includes solvation/desolvation and entropy in addition to the other enthalpic terms.

### Statistics

Unpaired two-tailed t tests or one way ANOVA were performed on triplicate or more data sets using GraphPad Prism 3.0 (Graphpad Software, inc.). *P* -values are indicated as *, <0.05; **, <0.01; and ***, <0.001.

## RESULTS

### PP2A functions downstream of ATM to regulate AKT

In a previous study from our laboratory, we demonstrated that ATM indirectly regulates the phosphorylation of AKT at S473 via an okadaic acid (OA) sensitive phosphatase (21). It is known that several oncogenic DNA viruses such as SV40 and Polyoma interfere with PP2A activity in infected cells with SV40 small t-antigen and PyMT blocking PP2A activity by replacing the B subunit (64–66) (**Supplemental Fig.1A**). After examining the effect of an ATM kinase inhibitor (ATMi) in a panel of tumor cell lines including human U87 and U1242 glioma cells as well as HEK293 and HEK293T carcinoma cells, inhibition of AKT phosphorylation was observed across the board relative to untreated cells with the exception of HEK293T cells (**Supplemental Fig.1B**). HEK293T expresses SV40 t/T antigens, which inversely correlates with the lack of pAKT inhibition by ATMi, suggesting similar or overlapping mechanism of action. To build on this finding, we infected U87 and U1242 cells with retroviruses expressing PyMT or not (**Supplemental Fig. 1C, D**). ATMi did not appear to reduce the levels of pAKT in glioma cells overexpressing PyMT. This effect was independent of exposure to ionizing radiation (IR) (21, 67). The presence of the PyMT gene in these cell populations was verified by PCR (**Supplemental Fig. 1E**). We confirmed the ability of OA to increase the levels of phosphorylated AKT (21), this time in a nuclear extract with nM concentrations suggesting that PP2A is acting on AKT in the nucleus (**Supplemental Fig. 1F**), perhaps as a complex in direct physical association with AKT (68). Altogether, these results suggest that ATM regulates the phosphorylation of AKT indirectly via the inhibition of PP2A activity.

### Generation of PR65 KO MEFs complemented with S401 mutant alleles

Ruediger *et.al*. demonstrated that PR65 is an essential gene, since knocking out PR65 resulted in embryonic lethality (36). We generated homozygous PR65 conditional knockout (CKO) mouse embryonic fibroblasts (MEFs) after breeding heterozygous 129S-*PPP2R1A^tm1wltr/j^* mice to obtain 129S-*PPP2R1A^tm1wltr/tm1wltr^* (CKO/CKO) mice. Floxing out PR65 exon 5 and 6 generates PR65 KO MEFs (**Fig. 1A**). MEFs generated from 129S-*PPP2R1A^tm1wltr/tm1wltr^* mouse embryos (37) were immortalized by passaging once per week and infecting the cells with a lentivirus expressing human TERT. We generated two site-specific mutations at the S401 codon of PR65 using the retroviral vector pMIG-Aalpha WT as template (32): a S401A mutant that cannot be phosphorylated and a S401D phospho-mimetic mutant. The MEFs were then separately infected with the retroviruses expressing Flag - tagged PR65 WT, S401A or a S401D mutant in addition to GFP (via an IRES). The endogenous PR65 alleles were then floxed out by infecting the cells with an adenovirus expressing Cre recombinase (Ad-Cre). Complete floxing was confirmed by PCR (**Fig. 1B**). MEFs that were not complemented with virus expressing PR65 did not survive after infection with Ad-Cre (**Fig. 1C**), in line with that PR65 is essential in the mouse (36). Pooled MEFs from each infection were shown to be close to 100% GFP+ (**Fig. 1D**). Western blotting with anti-PR65 and -Flag antibodies showed that at the protein level all endogenous PR65 (**Fig. 1E**, left panel; lower band), was replaced with the slightly larger Flag-PR65 protein expressed from the retrovirus. Altogether, we generated a panel of isogenic WT, S401A, and S401D cell populations expressing PR65 at approximately the same levels as endogenous PR65 (compare band intensity of PR65 doublet (**Fig. 1E**, right panel; lane 1).

### ATM kinase phosphorylates PR65 *in vitro*

The PR65 S401 residue is part of an – S/Q-ATM kinase signature motif highly conserved in PR65’s among mammals including human and mouse and is a *bona fide* substrate for ATM phosphorylation (31). To obtain an abundant source of co-expressed ATM and PR65 for further analysis and experimentation, we first attempted to isolate stably transfected HEK293 cells co-expressing mRuby2-PR65 and YFP-ATM after selection with G418 (expressed from YFP-ATM) and isolating cells expressing both YFP and Ruby. However, we were only able to recover YFP-ATM/S401A cells (**Supplemental Fig. 2A**), suggesting that over-expression of ATM with either S401D or WT was not tolerated.

**Figure 2.**
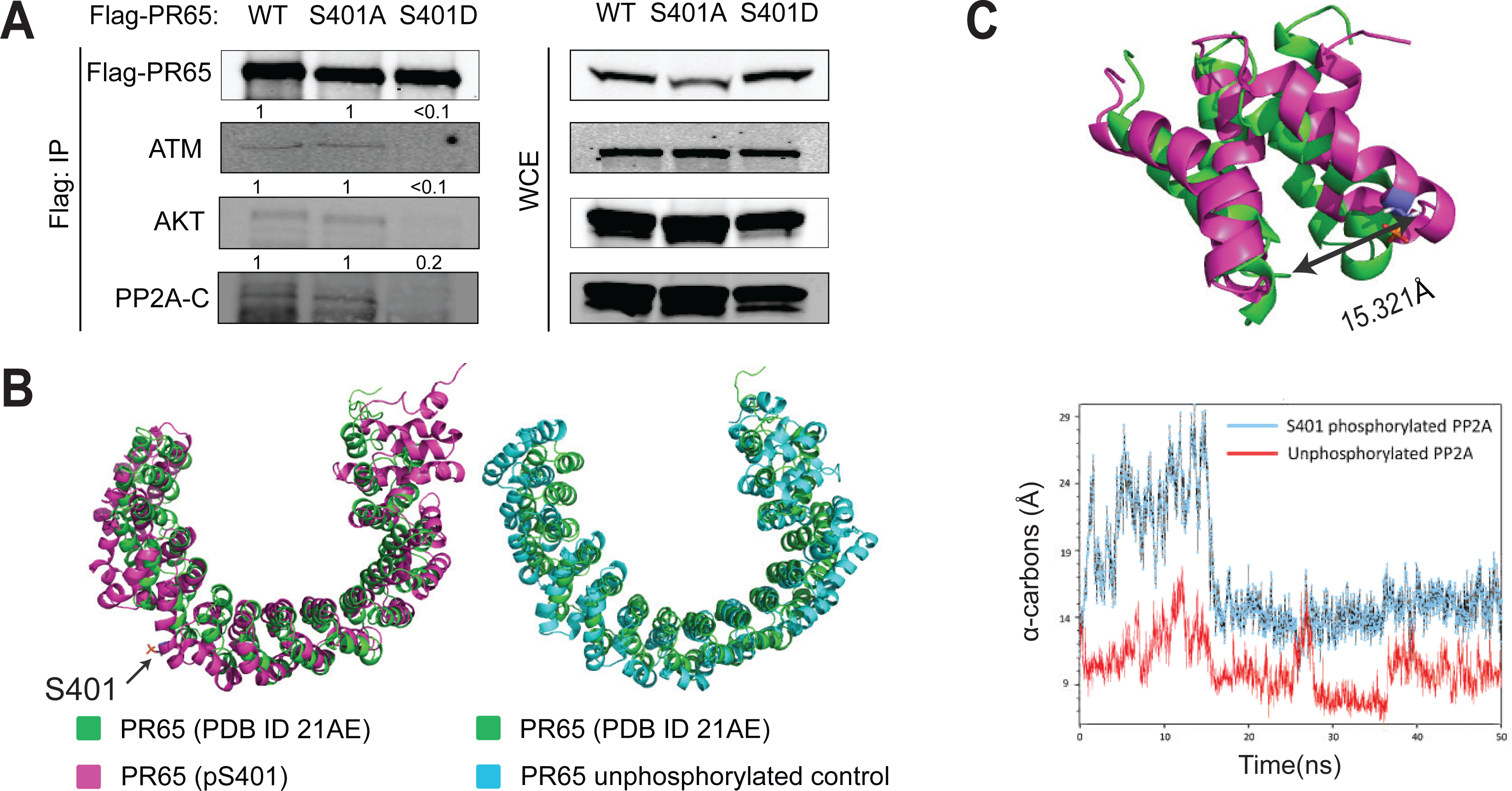
Phospho-mimetic S401D prevents the association of PR65 with ATM, AKT, and PP2A catalytic subunit. **(A)** Whole cell lysates of HEK293 cells stably expressing Flag-PR65 (WT, S401A and S401D) were immunoprecipitated with anti-Flag beads. Immunoprecipitates (*left panel*) and whole cell extracts (*right panel*) were separated on an SDS-PAGE gel and analyzed by western blotting using anti-Flag, anti-ATM, anti-AKT and anti-PP2A-C antibodies. **(B)** MD simulations suggests that a conformational change occur when S401 is phosphorylated. (*Left*) Top view of the *in silico* S401 phosphorylated PR65, after 50 ns, aligned to the original experimental structure (PDB ID 2IAE) with a root-mean square deviation (RMSD) of 8.05 Å. PR65 (p-S401) is shown in magenta, with p-S401 represented as stick figures. PR65 from protein database (PDB ID 21AE) is shown in green. (*Right*) Top view of the unphosphorylated control structure of PR65 subunit, after 50 ns, aligned to the original experimental structure taken from PDB ID 2IAE (RMSD = 6.51 Å). Unphosphorylated control PR65 is shown in cyan, and the original experimental structure shown in green. Molecular modeling studies also indicate that phosphorylation at S401 could cause a conformational change which could affect its association with a binding partner. **(C)** Two 50 ns molecular dynamics simulations were carried out using the S401 phosphorylated PR65 chain as well as the unphosphorylated structure. Comparing only the residues surrounding S401 (residues 375-454), the RMSD of the phosphorylated subunit and the unphosphorylated experimental structure was found to be 4.05 Å while the RMSD of the control and the unphosphorylated experimental structure was found to be 2.55 Å. At the end of the simulation, the distance between α-carbons for residues S401 and A431 for the unphosphorylated original structure, the S401 phosphorylated structure, and control structure were 10.873 Å, 15.321 Å and 9.498 Å, respectively (*top panel*). Initially, there was a readjustment phase in the phosphorylated MD simulation in the region surrounding the phosphorylated S401 residue after which it stabilized at 18 ns (*bottom panel*). Altogether this result suggests that phosphorylation of PR65-S401 could affect the interaction with its binding partners and result in the dissociation of the holoenzyme

To determine whether ATM phosphorylates PR65 *in vitro* and to confirm previously documented results (31), YFP-ATM was immunoprecipitated from HEK293 cells expressing YFP-ATM only (**Supplemental Fig. 2B**) and utilized for *in vitro* kinase assays with [γ-^32^P] ATP and GST-PR65FL as substrate. The result shows that ATM kinase phosphorylates PR65 *in vitro* and that phosphorylation is significantly reduced in the presence of an ATMi (**Supplemental Fig. 2C**, compare lanes 5 and 6). Similarly, GST-p531-100, a known substrate of ATM kinase at p53 S15 (40), was also phosphorylated and inhibited in the presence of the ATMi. This result confirms that ATM kinase is able to phosphorylate PR65 *in vitro* in line with a previous report (31).

### A phospho-mimetic (S401D) mutant prevents the association with ATM, PP2A catalytic subunit and AKT

Goodarzi et al. showed that PP2A exerts its control over ATM by interacting via the PR65 subunit in a manner lost in irradiated cells (25). To determine whether the S401D amino acid replacement affected the interaction with ATM, we immunoprecipitated over-expressed Flag-PR65 WT, -S401A and -S401D, respectively, from transduced HEK293 cells. Flag-PR65 IP’s were then analyzed for the presence of endogenous ATM, the catalytic subunit of PP2A (PP2A-C), and AKT. We found an association of ATM, PP2A-C, and AKT with the WT and the non-phosphorylatable S401A but not with S401D (**Fig. 2A**). The lack of an interaction between S401D and ATM but not WT or S401A suggests that the PP2A Core enzyme (AC) is possibly in a complex with ATM, in agreement with previous work (25). That the S401D phospho-mimetic disrupts this complex is new information and fits well with the dissociation of ATM and PP2A complex after irradiation and the phosphorylation of PR65 S401 by ATM (25, 31). That AKT is found complexed with PR65 is no surprise (68), but that S401D disrupts this interaction is intriguing, even though we cannot tell from this result whether ATM and AKT are in the same or separate PR65 complexes.

To support a critical role for S401 phosphorylation by ATM in the disruption of such physical interaction we performed molecular modeling studies based on the known PR65 crystal structure (PDB ID 2IAE). Two 50 ns Molecular Dynamics runs were carried out with phosphorylated S401 PR65 as well as the unphosphorylated structure (**Fig. 2B, C**). The result suggests that phosphorylation at S401 induces a local movement of the α-helices on either side of the helix (residues 375-454) encompassing S401 ultimately causing a conformational change of the entire PR65 protein, particularly at its N-terminal portion. This conformational change results in the distortion of the horseshoe shape of the subunit by *∼*4.5 Å compared to the unphosphorylated structure (15.321-10.873). Altogether, this modeling suggests that phosphorylation at S401 could cause the dissociation of the physical interaction with its binding partners ATM and AKT.

### Growth and radiation signaling are altered in S401 mutant cells

We then examined whether AKT signaling and growth was affected in response to insulin stimulation and radiation (21). WT, S401D, and S401A MEFs were serum-starved prior to stimulation with insulin followed by western blotting for pAKT (S473) (**Fig. 3**). We found that S401D cells responded more robustly to insulin when pAKT levels were determined over the course of 60 minutes (**Fig. 3A**), which correlated with an elevated growth rate of S401D relative to WT and S401A cells (**Fig. 3B**). Similar to AKT signaling, ERK signaling is associated with pro-survival and increased cell growth after low radiation doses, which in part is regulated by ATM (45, 69). When we examined ERK signaling in unstimulated cells we observed elevated pERK1/2 levels in S401A and more so in S401D cells compared with WT, suggesting that AKT and ERK signaling are positively affected when S401 is mutated, with S401D cells being affected the most (**Fig. 3C**). Furthermore, after a single radiation dose of 2 Gy, S401 mutant cells showed dampened responses when analyzed for both γ-H2AX and pAKT by western blotting (**Fig. 3D**). It is well-established that ATM phosphorylates H2AX at S139 (γ-H2AX) in response to radiation at the sites of DSBs (70). Furthermore, PP2A modulates DSB repair via the dephosphorylation of γ-H2AX to reverse chromatin accessibility (30). On examining the γ-H2AX levels in more detail, we found that in WT cells the levels of γ-H2AX peaked at 15-30 min post irradiation and then declined. However, H2AX phosphorylation was reduced in both S401A and S401D cells with the latter trending even higher than WT and S401A at 60 minutes (**Fig. 3D**). Similarly, pAKT levels increased after IR in WT and less so in S401A and S401D cells. These results suggest that after low dose radiation, S401 mutant cells have dampened pAKT and γ-H2AX levels likely as a result of malfunctioning DDR. Additionally, S401D seems to overcome this initial blockade with a delayed response and increased γ-H2AX levels at later times. Altogether, we conclude that S401 mutant cells have aberrant responses to insulin and radiation yet distinct from each other, resulting in altered DDR signaling and proliferative activity compared to WT cells.

**Figure 3.**
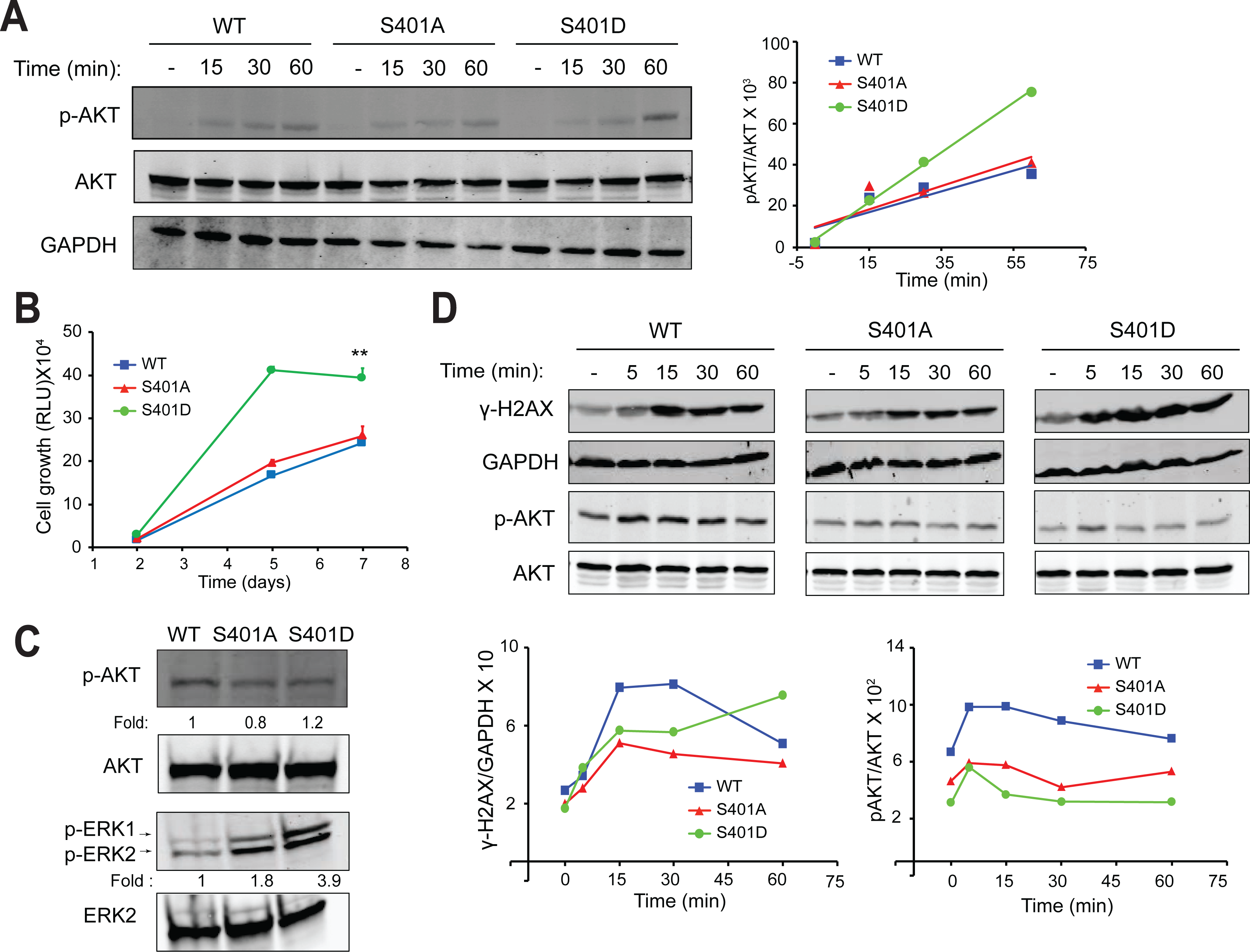
Growth and irradiation signaling is impaired in PR65 S401 mutant cells. **(A)** AKT phosphorylation is increased in S401D cells after insulin stimulation. MEFs (WT, S401A and S401D) were serum starved for 16 hours and then treated with insulin for 15-, 30- or 60-min. Whole cell extracts were separated on an SDS-PAGE gel and analyzed by western blotting with anti-pAKT (S473), anti-AKT, anti-GAPDH (loading control) antibodies (*left panel*). Western blots were quantified using Image Studio Lite. p-AKT (S473) levels were normalized to total AKT protein levels (*right panel*). **(B)** S401D cells have increased growth rate relative to S401A and WT. MEFs were grown at different dilutions in 96-well plates. Cells were analyzed by Cell titer-Glo® at 2, 5, and 7 days to determine growth. Data points; Growth, Relative Luminescence Units (RLU). Error bars; mean ± SEM (n = 3). Statistical analysis was carried out using unpaired, two-tailed *t*-tests. *P*-values expressed as *(*P* < 0.05) and **(p < 0.01) were considered significant. At 7 days, WT vs. S401D; p=0.0026, S401A vs. S401D; p= 0.0118. **(C)** ERK phosphorylation is elevated in S401D cells. Whole cell extracts of MEFs were separated on an SDS-PAGE gel and analyzed by western blotting using anti-pAKT, anti-AKT, anti-pERK and anti-ERK antibodies. pAKT (S473) levels were normalized to total AKT protein levels and anti-pERK (T202/Y204) levels were normalized to total ERK levels. Increased pERK levels correlate with increased growth of S401D cells. **(D)** S401 cells show dampened response to irradiation. MEFs were exposed to 2 Gy of ionizing radiation and collected after 5, 15, 30 and 60 minutes. Whole cell extracts were separated on an SDS-PAGE gel and analyzed by western blotting using anti-pAKT, anti-AKT, anti-γ-H2AX, and anti-GAPDH antibodies (*top panel*). Western blots were quantified using Image Studio Lite (Li-Cor). p-AKT levels were normalized to total AKT protein levels and γ-H2AX levels were normalized to GAPDH (*bottom panels*).

### PR65 mutants have impaired G2/M checkpoint resulting in aberrant mitosis and elevated levels of mitotic catastrophe

PP2A plays distinct roles at different stages of mitosis while associated with different B subunits (71, 72). PP2A regulates mitotic entry and exit, as well as playing a role in the protection of centromeres by inhibiting PLK1 and Aurora A kinases (73–77). Taking into account the importance of PP2A in mitosis at multiple steps and elsewhere during the cell cycle, we examined whether S401 mutant cells show any perturbations in mitosis and cell cycle checkpoints. We found that the mitotic index (cells undergoing mitosis/total number of cells) was reduced in WT cells after radiation but not in S401 mutant cells, suggesting that the radiation-induced G2/M check point is intact in WT cells, as expected, but not functioning properly in the mutants (**Fig. 4A, B**). Furthermore, S401 mutants had significantly increased numbers of basal and radiation-induced (2-3 fold) chromosomal aberrations (**Fig. 4C**). To support this finding, live cell imaging of MEFs expressing a histone H2B-mCherry reporter revealed that the S401 mutant cells exhibited abnormal mitoses including the formation of micronuclei and elevated levels of mitotic catastrophe, and the formation of tetraploid cells whose frequency increased after exposure to radiation (**Fig. 4D**, left panel). Again, S401D cells showed more aberrations than S401A. Furthermore, in undamaged cells, the length of mitosis in S401 mutant cells was significantly longer than in WT cells (**Fig. 4D**, right panel). To examine mitosis in more detail, we then focused on exit from mitosis. PLK1 is known to be critical during G2/M entry as well as for successful chromosome separation and exit from mitosis (78). Normally, pPLK1 (T210) is elevated in mitosis and upon exit is dephosphorylated by PP2A (79). A nocodazole blockade synchronized the cell panel in mitosis and following washout, we collected cells for western blotting with anti-pPLK1 (T210) antibody (**Fig. 4E**). Mitotic S401 mutant cells showed elevated pPLK1 levels compared with WT at 1 hour and exited mitosis faster than WT cells with S401D showing the most pronounced effect (**Fig. 4F, G**). Together, both mitotic entry and exit are abnormal in S401 mutant cells reflected in significantly longer mitosis and clear signs of struggle to get through mitosis.

**Figure 4.**
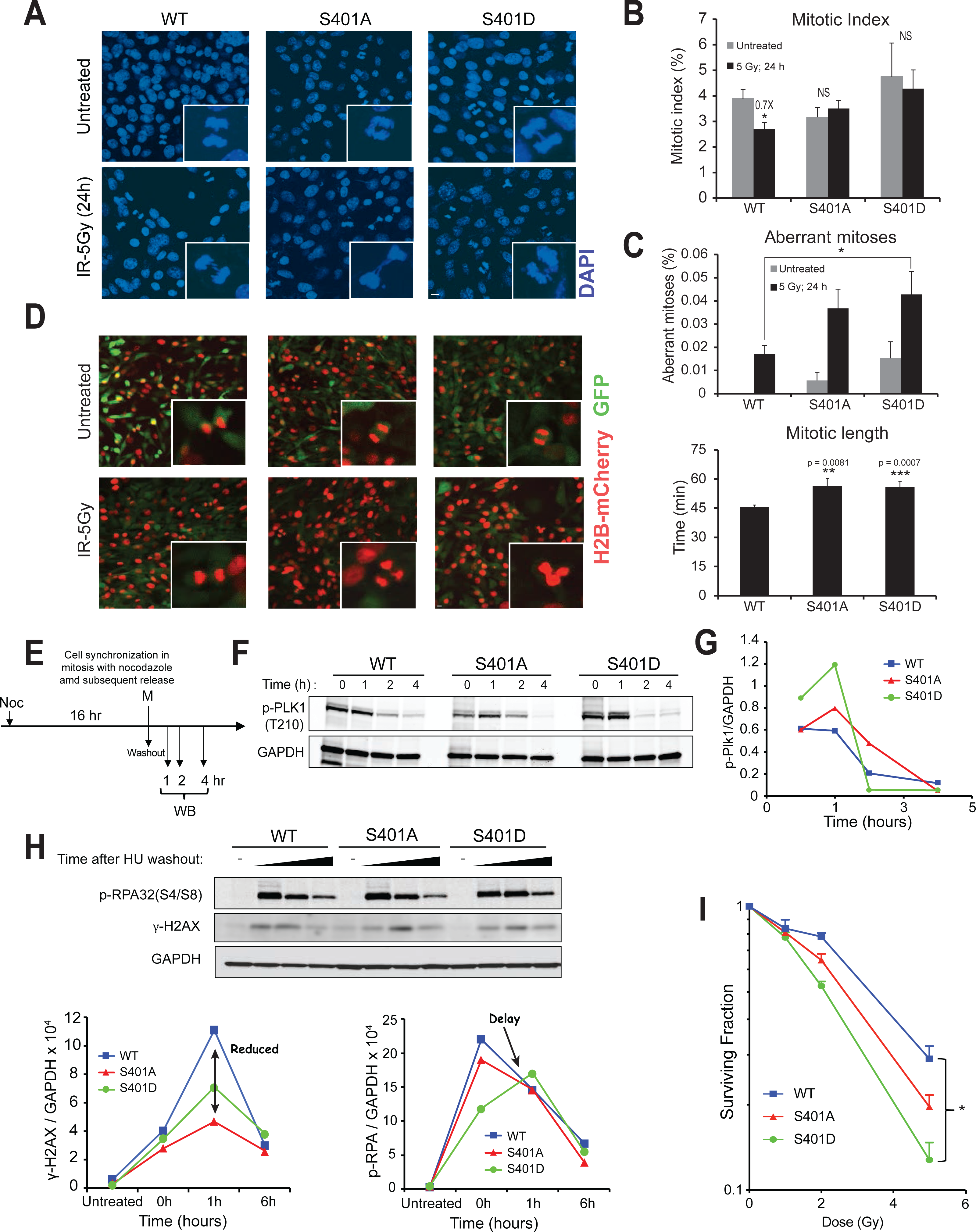
PR65 mutants accumulate mitotic aberrations leading to mitotic catastrophe and radiosensitization. (**A, B and C**) S401 mutant cells have defective G2/M checkpoint. MEFs were exposed to 5 Gy of ionizing radiation, fixed after 24 hours and stained with DAPI to label nuclei. Mitotic cells and total number of cells were counted. The mitotic index was calculated by dividing the number of cells undergoing mitosis in a population to the number of cells not undergoing mitosis. Five separate fields were assessed for each group Scale bar is 10 μm. (**B**). Mitotic indices were quantified from **A**. Error bars; mean ± SEM Statistical analysis was carried out using unpaired, two-tailed *t* - tests. *P* values expressed as *(*P* < 0.05) were considered significant; *n.s*. not significant (*P* > 0.05). WT: Untreated vs IR; p = 0.0251. (**C**) Aberrant mitoses were quantified from **A**. Five separate fields were assessed for each group. Error bars; mean ± SEM. Statistical analysis was carried out using unpaired, two-tailed *t* - tests. *P* values expressed as *(*P* < 0.05) were considered significant; *n.s*. not significant (*P* > 0.05). IR: WT vs S401D; p = 0.0228 (**D**) MEFs were infected with a H2B-mCherry virus construct to monitor chromatin structure in mitosis. MEFs were exposed to 5 Gy and subjected to live-cell video imaging for 8 hours on the Zeiss Cell Observer Spinning Disc confocal microscope. Representative still images (*left panel*) and quantified mitotic length (*right panel*) Scale bar is 10 μm. **E**) S401D cells quickly exit mitosis after mitotic synchronization. Cells were arrested in M phase with 16 hours of nocodazole treatment. Mitotic cells were collected by shake-off, reseeded in complete media, and collected after 1, 2 and 4 hours for western blotting with anti-pPLK1 (T210) antibody, a marker of mitotic exit (**F**), and plotted (**G**). Faster pPLK1 dephosphorylation occurs in S401D cells expected to speed up mitotic exit. Altered PP2A-S401D target specificity/activity might explain quicker exit from mitosis. (**H**) S401D cells show delayed replication restart. MEFs were treated with 1 mM of hydroxyurea (HU) for 2 hours followed by drug wash-out and collected at 0, 1, and 6 hours after wash-out. Western blotting with antibodies specific for pRPA (S4/S8) and γ-H2AX normalized to total GAPDH. (**I)** S401 mutant cells are more radiosensitive than wild-type MEFs. MEFs were exposed to 1, 2 or 5 Gy and radiosurvival determined at 10 days. Surviving fractions were determined by crystal violet staining and colony counting. Data points, surviving cells plotted as fraction of control (-irradiation). Results are presented as mean ± SEM, (n = 4). Statistical analysis was carried out with longitudinal ANOVA on the survival fractions. *P*-values expressed as *(*P* < 0.05) were considered significant. WT vs. D, p = 0.0302.

To get better insight into what occurs during S-phase we then examined the recovery of collapsed replication forks in the cell panel by determining pRPA32 (S4/8) and γ-H2AX levels after the release from a hydroxyurea (HU) blockade (**Fig. 4H**). A previous study showed that PP2A is critical for the dephosphorylation of pRPA32, cell cycle reentry and efficient DSB repair presumably by HRR (80). We found that S401 mutant cells had reduced γ-H2AX levels after HU release, which peaked at *∼*1 hour post washout (**Fig. 4H**; bottom left panel). Perhaps more importantly, S401D cells showed a delay in the removal of pRPA32 (**Fig. 4H**; bottom right panel) indicative of an alteration in PP2A activity during fork recovery. This result suggests that the recovery of collapsed replication forks is impaired in S401 mutant cells and that PP2A, at least in S401D cells, might adversely affect pRPA32 dephosphorylation and perhaps subsequently impact HRR. In addition, a colony-forming radiosurvival experiment demonstrated that S401D cells were also significantly more radiosensitive than WT cells, with S401A trending more radiosensitive (**Fig. 4I**). The same pattern of sensitivity was seen when the cell panel was exposed to camptothecin (CPT), a topoisomerase I inhibitor, known to produce DSBs preferentially in S and G2 (**Supplemental Fig. 3**).

Altogether, increased chromosomal aberrations seen in untreated S401 mutant cells (and more so after radiation) are suggestive of impaired G2/M checkpoint and aberrant mitosis (entry, duration, and exit), resulting in elevated levels of mitotic catastrophe and increased sensitivity to DNA damage. In addition, replication fork recovery and DSB repair were impaired, especially in the S401D cells. It was however unexpected to see that the impact on survival was relatively minor considering the major negative effects on the G2/M checkpoint and increased mitotic struggle seen in S401 mutant cells.

### S401 mutants have aberrant DSB repair

In addition to reversing changes in chromatin structure caused by H2AX phosphorylation, PP2A is believed to facilitate NHEJ by dephosphorylating KU70/80 and DNA-PKcs, which are critical participants in NHEJ (14), and to promote their release from DNA (81). Furthermore, PP2A restores kinase activity of phosphorylated, inactive DNA-PKcs in vitro to regulate NHEJ (82). To determine how S401 alterations might affect DSB repair we then carried out *in vivo* as well as *in vitro* end joining assays. First, in a reporter assay, WT, S401A and S401D mutant cells were transfected with a linearized pCSCMV:tdTomato plasmid with incompatible restriction endonuclease DNA ends unable to ligate. Once the plasmid is repaired and circularized, the red fluorescence protein (RFP) expressed is a measure of end joining activity (**Fig. 5A**). Post transfection, cells were collected at various times and RFP+ cells quantified. At 16 hours the S401A cells had almost twice the number of RFP+ cells relative to WT cells whereas the S401D cells had much less than the WT cells throughout (**Fig. 5B**, top panel). To examine early repair, we quantified the circularized plasmid at 1 and 4 hours after transfection and determined end joining yield by quantitative PCR (**Fig. 5B**, bottom panel). The PCR result is in agreement with what we observed with the reporter assay, i.e., S401D cells have compromised end joining. S401A cells probably did not have enough time to translate into elevated repair over WT seen at 16 hours.

**Figure 5.**
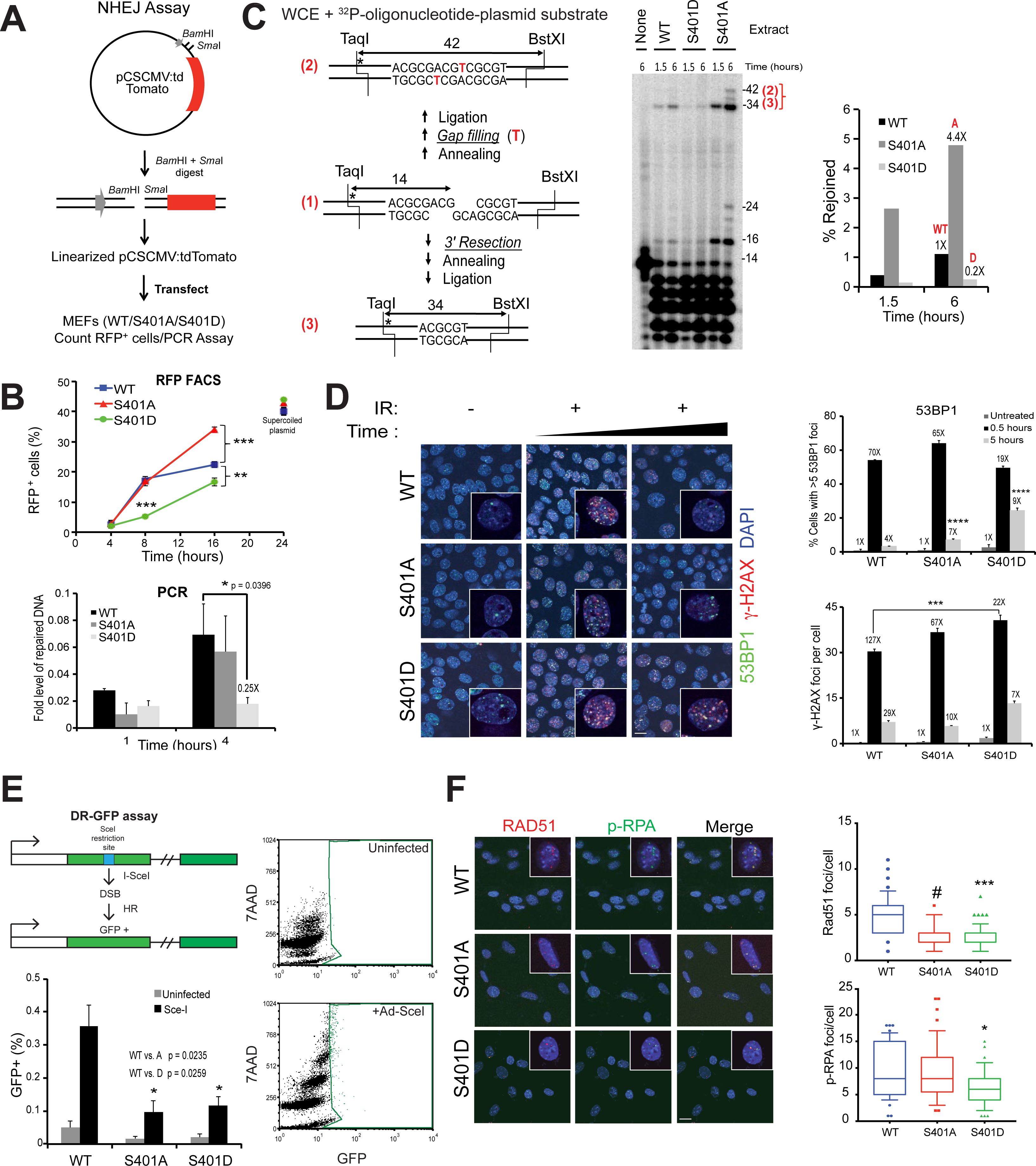
DSB repair is abnormally high in S401A cells while being suppressed in S401D. (**A**) Scheme for *in vivo* end joining (EJ) assay. Linearized Tomato expression plasmid was transfected into MEFs followed by quantification of RFP+ cell (Nexcelom Cellometer) and PCR. (**B**) Quantification of EJ by RFP flow cytometry (*top panel*). S401A cells have super-active EJ at later time point after transfection whereas S401D cells have suppressed EJ throughout. qPCR assay was carried out to determine early EJ (*bottom panel*). EJ in S401D cells was suppressed by 75% at 4 hours in line with the results by RFP flow. **(C)** *In vitro* end joining. End joining of an internally labeled (*) plasmid substrate with partially complementary ends in MEF extracts (*left panel*). Following incubation for the indicated times, substrates were cut with BstXI and TaqI and analyzed on sequencing gels (*middle panel*). Gap-filling products (42-base fragment) and resection-based products (34-base fragment) were quantified. The 24- and 16-base fragments are the corresponding head-to-head end joining products. Graph shows sum of the major 34-base fragments in each case (*right panel*). (**D**) Mutant S401 cells show aberrant IR-induced foci with delayed disappearance of 53BP1 and γ-H2AX foci in S401 mutant cells (*left panel*). MEFs were seeded in a chambered slide. Cells were exposed to 2 Gy of ionizing irradiation and fixed after 0.5 and 5 hours. Cells were immuno-stained with anti-53BP1 and anti-γ-H2AX and counterstained with DAPI (blue) to label nuclei. Representative images 53BP1 (Green), γ-H2AX (red) and DAPI (blue). Images were acquired on Zeiss LSM 710 confocal microscope at 63x power. Repair foci remain significantly longer in S401A and -D (esp. D) than in WT cells suggesting aberrant DSB repair. Delayed disappearance of 53BP1 (*top, right panel*) and γ-H2AX (*bottom, right panel*) foci in S401 cells post irradiation. Error bars; mean ± SEM. Statistical analysis was carried out using unpaired, two-tailed *t* - tests. p*-*values are shown as ***(p < 0.005) is considered significant. Scale bar is 10 μm. (**E**) S401A and D cells are impaired in homologous recombination using the DR-GFP assay (*top, left panel*). PR65-DR-GFP (WT, S401A and S401D) cells were infected with Ad-SceI (*bottom, right panel*) or not (*top, right panel*) and 72 hours later, cell populations analyzed by GFP flow cytometry after incubation in 0.1% Triton-X-100 in PBS. Results (*bottom, left panel*) are presented as mean ± SEM (30,000 events), n = 3, *p < 0.05 relative to WT Scale bar is 10 μm. **(F)** Reduced Rad51 and pRPA foci in S401 mutants post-CPT treatment. MEFs were treated with 100 nM of CPT for 6 hours, fixed and immunostained with anti-pRPA32(S4/S8), anti-Rad51 antibodies, and counterstained with DAPI (blue). Representative images Rad51 (Red), pRPA (Green) and DAPI (blue) images acquired on a Zeiss LSM 710 confocal microscope at 63x power (*left panel*). Quantification of Rad51 and pRPA32 (S4/8) foci (*right panels*). Rad51 foci per cell were counted in 5 separate fields. Error bars; mean ± SEM. Statistical analysis was carried out using unpaired, two-tailed *t*-tests. *P*-values expressed as ***(p < 0.005) and ^#^(p < 0.001) were considered significant. For Rad51 foci, WT vs. S401A; p=0.0008, WT vs. S401D; p=0.0011. For pRPA foci, WT vs. S401D; p=0.0443.

To assess the possible effect of PP2A-mediated dephosphorylation on end joining *in vitro*, a ^32^P-oligonucleotide ligated into a plasmid substrate bearing partially complementary 5′ overhangs was incubated in whole-cell extracts from WT, S401A, and S401D cells for 1.5 or 6 hours (**Fig. 5C**). As in our previous work (43, 83), the predominant 34-base head-to-tail product as well as the 16-base head-to-head product reflects 3′ resection and annealing at a 4-base micro-homology (**Fig. 5C**, left panel). We observed dramatically reduced end joining activity with the extract from S401D, whereas the S401A extract markedly enhanced it and yielded an additional 42-base product corresponding to gap filling without resection (**Fig. 5C**, middle and right panels). Thus, while the critical PP2A dephosphorylation targets are currently unknown these results show that using S401 mutants as surrogates for mimicking (de)phosphorylated S401 is important for end joining by reporter as well as in cell-free in vitro assays. In line with these results, S401D cells showed abnormal, extensive and more persistent 53BP1 foci after radiation compared with WT (**Fig. 5D**). 53BP1 is critical for directing DSB repair towards NHEJ at the expense of HRR (84). 53BP1 foci co-localized with γ-H2AX, suggesting that NHEJ is significantly altered and delayed in S401D cells. Furthermore, S401A cells had fewer but more pronounced foci at 5 hours than either WT or S401D (**Fig. 5D**). Altogether, S401 mutant cells have significantly altered end joining activity in quality as well as temporal execution yet S401D and S401A differed in that the latter showed super-active end joining whereas S401D was severely compromised.

### S401 mutants are HRR impaired

Since NHEJ and HRR are in competition in S/G2 of the cell cycle and we observed irregular end joining in the S401 mutants, we then examined HRR in the cell panel. The PP2A holoenzyme associated with the B55 subunit is implicated in HRR via the modulation of ATM phosphorylation and the G1/S checkpoint (85), as well as the dephosphorylation of pRPA32 mediated by PP2A after replication fork recovery, which might impact HRR (80). To determine the importance of the S401 residue in HRR, we generated CKO/CKO MEFs with the DR-GFP repair reporter (44) followed by infection with either WT, S401A and S401D retrovirus and floxing of the endogenous PR65 gene with Ad-Cre (**Supplemental Fig. 4**) as before (see **Fig. 1B, E**). We saw that HRR was significant reduced in S401A and S401D cells represented by lower GFP+ events (**Fig. 5E**). During HRR, single stranded DNA generated during DNA resection is first coated with RPA and then replaced by RAD51 during Holliday junction formation and resolution (14). This step is essential for BRCA1 recruitment and efficient HRR (86). Therefore, we treated the cells with CPT and then co-stained for pRPA32 (S4/8) and RAD51. Both S401A and S401D cells had *∼*40% less RAD51 and pRPA32 foci compared with WT cells (**Fig. 5F**). This finding supports the result with the DR-GFP reporter and the conclusion that HRR is impaired in both S401A and S401D cells relative to WT. Altogether, it is quite clear that S401 mutant cells have rewired DDR and show aberrant DSB repair. Surprisingly, however, these mutants display fairly minor effects on cell survival after radiation and CPT treatment. This is particularly striking with the S401D cells since they are compromised in end joining as well as HRR.

### Increased aberrant chromosomal DSB repair and translocation in S401 mutants

Alternative end joining (Alt-EJ, now referred to as Theta-mediated end joining; TMEJ) and other micro-homology repair pathways are believed to become engaged when c-NHEJ (classical end joining) and/or HRR are not fully functional or being disengaged (46,47,87). However, TMEJ is highly mutagenic because of the promiscuous activity imposed by DNA polymerase Theta (Pol θ) - mediated chromosomal repair and rearrangements through short (2-4 bp) stretches of micro-homology in resected DNA ends. To assess TMEJ in our PR65 cell panel we examined DSB repair quality after CRISPR/Cas9-mediated DNA cleavage at the Rosa26 locus and by chromosomal translocation between the Rosa26 and H3fl3 loci on Chr. 6 and 11, respectively (46, 47). The fact that the Rosa26 cleavage site is positioned in the middle of an XbaI site allows for the elimination of any DNA resealing products without indels, which if present would destroy the XbaI site, as well as DNA that was not cut (**Fig. 6**). As expected, XbaI-resistant PCR fragments remained after cleavage with Cas9-Rosa26/H3f3b gRNAs expressed but not without (**Fig. 6A**).

**Figure 6.**
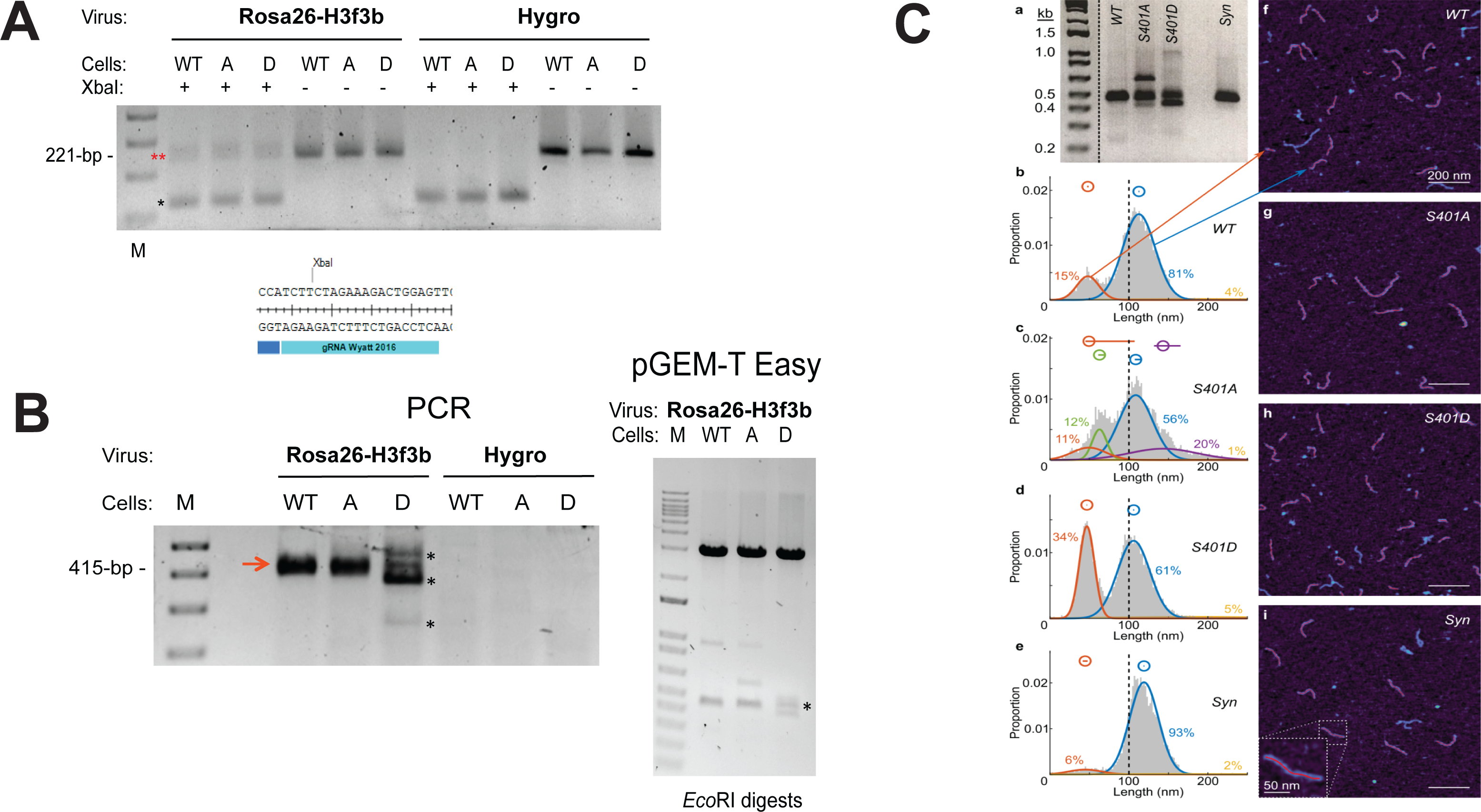
CRISPR-Cas9 mediated DSB repair. (**A**) **DSB repair at the Rosa26 (Chr 6) locus**. XbaI digestion removes background PCR DNA without indels (no NHEJ)* and leaves behind the *∼*221-bp PCR fragments with indels** (*top panel*). Primers: Chr6 Rosa26-F/Rosa26-R (47). The Rosa26 CRISPR-Cas9 gRNA target is spanning an XbaI-I restriction site (*bottom panel*). Thus, an indel would eliminate the Xba-I site. (**B**) Rosa26 (Chr 6) and H3f3b (Chr 11) translocation between DSBs generated by dual CRISPR-Cas9 cleavage. Primers: Chr6 Rosa26-F/Chr11 H3f3b-R. Major indels at translocation junctions are seen only in D cells* although translocation is seen in all three cell mixes (*left panel*). PCR fragments were cloned into pGEM-T Easy and sequenced (*right panel*). (**C**) Fragment size distributions of Rosa26-H3f3b translocation amplification products as determined by **(a)** 2% agarose gel electrophoresis and **(b)–(i)** high-speed atomic force microscopy (AFM). Four samples were amplified: wild-type (WT, *n* = 11,303 measured strands), S401A (*n* = 6,530), S401D (*n* = 12,160), and a synthetic fragment (Syn, *n* = 6,035) of equivalent length and sequence as WT. In (b)– (e), the normal probability density of amplicon lengths for each sample is shown as well as a Gaussian mixture model fit to each distribution. The probability density function of each single Gaussian fit is shown as a smooth colored curve, with that curve’s population proportion given in the same color. Also, above the probability density plots, the mean of each Gaussian fit is plotted as a hollow circle, and the 95% confidence interval of the mean is shown as a horizontal line (CI calculated from a bootstrap simulation with *n=*10,000) (**Supplemental Table 1**). In (f)–(i), high-speed AFM images of DNA amplicons from each sample are shown with 200 nm scale bars and lighter colors representing greater height. In frame (b), arrows point from the two main distributions to imaged strands in frame (f) that belong to the indicated population. In frame (i), a higher magnification of a single strand is shown. The red backbones shown on some amplicons represent the measured lengths of those amplicons; if an amplicon has no red backbone, it did not meet the image analysis algorithm’s quality standard and was not measured.

Because of the complexity of analysis and difficulties achieving meaningful statistical support for differences in the quality and quantity of end joining products between cells in the panel, we decided to examine the spectrum of translocations occurring between Chr. 6 and 11 after inducing dual chromosomal breaks (46, 47). TMEJ appears to play an important role in chromosomal translocations in human cells (88–91). However, in the mouse, the role of Pol Theta in translocation repair is more controversial (92).

Nevertheless, with PCR primers amplifying translocation events as an expected 415-bp fragment we found that the S401D cells produced relatively large insertions as well as deletions compared with either WT or S401A cells which gave bands closer to the expected PCR fragment size without large indels (**Fig. 6B**, left panel). The PCR fragments were cloned and some sequenced (**Supplemental Fig. 5**). Among the DNA sequences, we found indels with micro-homology in the DNA from S401D and S401A cells. Altogether, translocations generated in S401D cells are to a large extent associated with significant micro-homology end joining since predominant number of DNA sequences had short 2-4 bp repeats in the junction between Chr. 6 and 11. In line, with the results generated throughout this study using different DNA repair assays, we again noticed differences in the quality of DSB repair between the three cell populations.

### Assessing translocation events by atomic force microscopy

Because cloning of PCR fragments might be biased against isolating longer inserts many of which might have direct or inverted repeats (that could be unstable in bacteria), we decided to utilize a novel, high-throughput technique that does not require plasmid cloning. A recently developed alternative for determining translocations by indel size is by using atomic force microscopy (AFM) (51). High-speed AFM-based physical mapping of DNA allows single-molecule measurement of thousands of PCR products in a relatively short time. However, only the size of the indels are determined without establishing the DNA sequence but the assay is nevertheless a way to quantify differences in DNA repair quality between the three cell populations, simplify the process, and, importantly, allow for better statistical analysis. In this experiment, we noticed that S401A as well as S401D gave translocation indels seen on an agarose gel whereas few were seen from WT cells or a synthetic (*Syn*) template without any indel at the Rosa26-H3f3b junction that served as standard (**Fig. 6C**, panel a). On the other hand, when we examined the AFM results using the same PCR reactions as examined by agarose electrophoresis we noticed shifts in the size of indels between the three cell populations (**Fig. 6C**, panels b-e) and images of the DNA corresponding to the various curves (**Fig. 6C**, panels f-i). The *Syn* template produced sizes centered at 119 nm (blue; 93%) and a bulge of smaller sizes around 45 nm (orange; 6%) representing background noise products (**e, i, Supplemental Table 1**). WT cells predominantly gave sizes around 113 nm (blue; 81%) slightly smaller than with the *Syn* standard template, indicating smaller deletions at the junction as expected as well as smaller fragments around 48 nm (orange; 15%) some of which possibly represent deletions of 200-bp or more (**b, f**). The size pattern resulting from S401A was more complex with a main peak at 108 nm (blue; 56%), a broad peak centered at 143 nm and extending >260 nm (purple; 20%) in size, and a range of deletions with peaks between 49-61 nm (green; 12%, orange; 11%). Finally, S401D produced a predominant peak centered around 106 nm (blue; 61%), slightly smaller in size then either WT or S401A and an abundant peak centered around 47 nm (orange; 34%). The overall pattern and difference in fragment size distribution of indels seen between WT, S401A, and S401D by AFM are significant (**Supplemental Table 1**). The observed distinct peaks might represent repair events resulting from preferred microhomology hotspots such as repetitive DNA in resected ends that might serve as sinks for repair. Clearly, the quality of DSB repair is dramatically altered when S401 is replaced with either a null or a phospho-mimetic substitution reflecting the importance of PR65 S401 (de)phosphorylation during the DDR. However, to determine the mechanism behind these differences and whether it represents TMEJ or other (backup) DSB repair goes beyond the scope of this report and will have to wait for a more thorough future investigation.

### S401 (de)phosphorylation shuttles PR65 between the nucleus and the cytoplasm

To return to the overarching goal of this study and how growth, cell cycle regulation, and DSB repair could be so fundamentally altered by a single amino acid change mimicking the phosphorylation or not of PR65 at S401 by the ATM kinase, we wanted to explore this issue more thoroughly. In neurons, ATM phosphorylates PR65 at S401 resulting in nuclear-cytoplasmic shuttling of HDAC4 and PR65 (31). Thus, we examined the cellular location of WT, S401A and S401D PR65 in the MEFs using Flag antibody. In WT cells PR65 was present in the cytoplasm as well as the nuclear compartment whereas S401A was predominantly nuclear and in S401D exclusively cytoplasmic with little change after radiation except that in the WT and S401A cells PR65 appeared more in punctate nuclear foci (**Fig. 7**). After irradiation of WT cells, PR65 seemed to translocate to the cytoplasm even though it is unlikely that all PR65 complexes participate in a unison manner blurring the response and interpretation thereof. This observation in addition to the sequestration of S401D mutant in the cytoplasm suggests that the phosphorylation of PR65 at S401 causes it to translocate from the nucleus to the cytoplasm in line with a previous report (31). To explore the mechanism, we treated the cells with leptomycin B (LMB), which is a very specific inhibitor of nuclear export protein CRM1 (chromosome region maintenance 1) responsible for the nuclear export of many proteins and RNAs (93). After LMB treatment of WT cells, PR65 began to accumulate in the nucleus and combined with irradiation this effect was even more pronounced (**Fig. 7B**, compare middle panels at 30 min +/- LMB). In support of this finding, S401A cells did not respond to radiation with PR65 remaining in the nucleus whereas in S401D cells PR65 was almost exclusively cytoplasmic regardless of being irradiated or not (**Fig. 7A**). These results suggest that nuclear export of PR65 after radiation and phosphorylation by ATM kinase is mediated by CRM1 (**Fig. 7C**).

**Figure 7.**
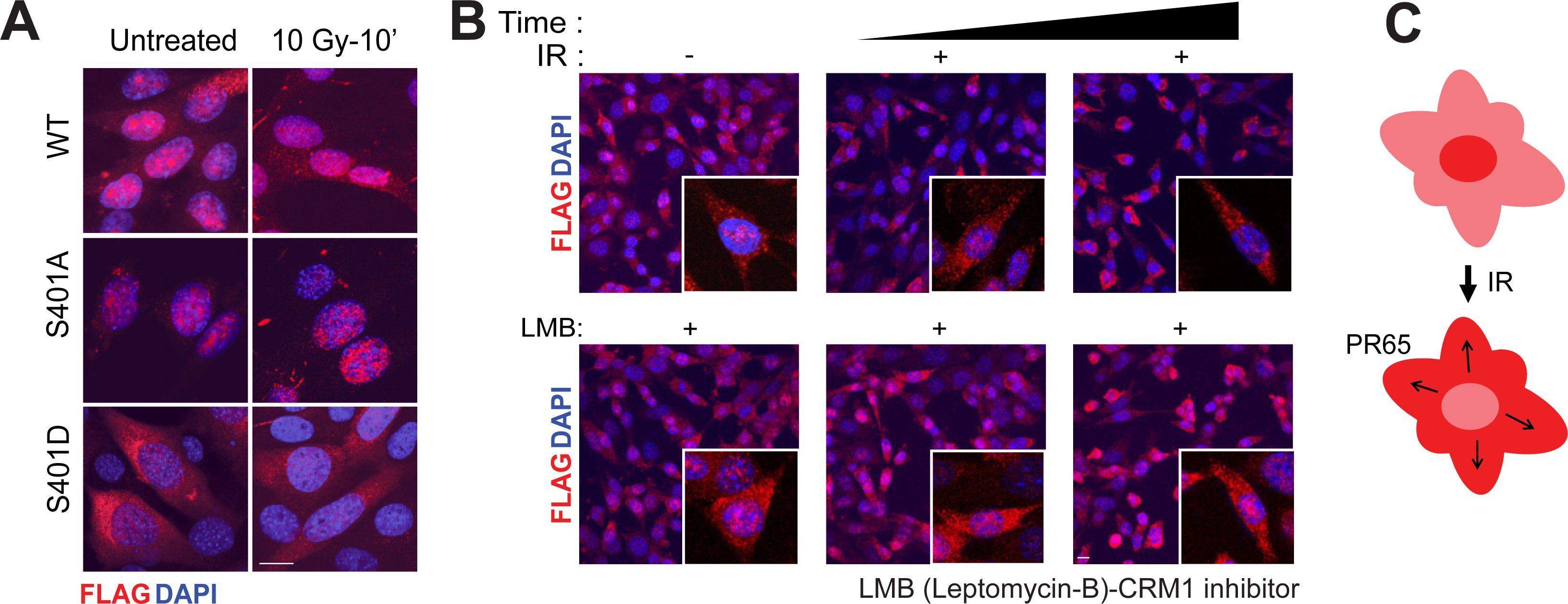
Radiation-induced nuclear export of WT PR65 is mimicked by S401D. (**A**) Wild-type MEFs grown in chamber slides were irradiated with 10 Gy with or without leptomycin B (LMB) present (1 ng/ml; -3 hour) in the medium. Cells were fixed at 30 and 60 minutes after irradiation and stained with anti-Flag antibody to detect PR65 (red) and counterstained with DAPI (blue). Images were acquired on Zeiss LSM 710 confocal microscope at 40x power. Irradiation results in cytoplasmic accumulation of Flag-PR65 seen best at 60 min (*top panel*). LMB blocks PR65 nuclear-cytoplasmic shuttling as shown best by the decreased nuclear and increased cytoplasmic accumulation at 30 min (*bottom panel*) Scale bar is 10 μm. (**B**) MEFs were grown on chamber slides and irradiated with 10 Gy, fixed at 10 min and processed as in (A). S401D markedly showed no nuclear staining prior to or post irradiation whereas both WT and S401A cells have Flag-PR65 nuclear and cytoplasmic staining Scale bar is 10 μm. (**C**) Model for radiation-induced nuclear export of PR65 mediated by S401 phosphorylation (see text).

### PR65 shuttles to the cytoplasm by CRM1 via a nuclear export sequence

After DNA damage several DDR proteins including p53, BRCA1, and BRCA2 are exported out of the nucleus through NES-mediated CRM1 processes (94–96). To determine whether PR65 might have a NES, we scanned the PR65 amino acid sequence for putative hydrophobic leucine residue motifs using the online tool NetNES1.1 (http://www.cbs.dtu.dk/services/NetNES/). We identified two leucine residues, L370 and L373, that scored positive as potential residues of a NES sequence between PR65 amino acids 365-373 (LLPLFL370AQL373) which resembles the consensus NES LxxxLxxLxL (97) (**Supplemental Fig. 6A**). Nuclear export of proteins harboring a functional NES sequence requires CRM1 binding to the target (98–100). We therefore examined whether CRM1 and PR65 interact by performing co-immunoprecipitation of lysates of HEK293 cells expressing Flag-CRM1 and mRuby2-PR65 also including a plasmid expressing the RanQ69L mutant that is incapable of exchanging GTP for GDP and prevent the dissociation of the Ran-CRM1 complex in whole cell extracts (101, 102). The immunoprecipitation/western blotting experiment shows that CRM1 interacts with PR65 (**Fig. 8A**). One prototype CRM1 interactive partner is Snurportin-1 (Exportin-1) for which a co-crystal structure has been determined (PDB ID 3GB8) This structure was then used for identifying possible CRM1-PR65 hydrophobic interactions by HINT analysis modeling hydrophobic protein interactions under aqueous condition (103–105).

**Figure 8.**
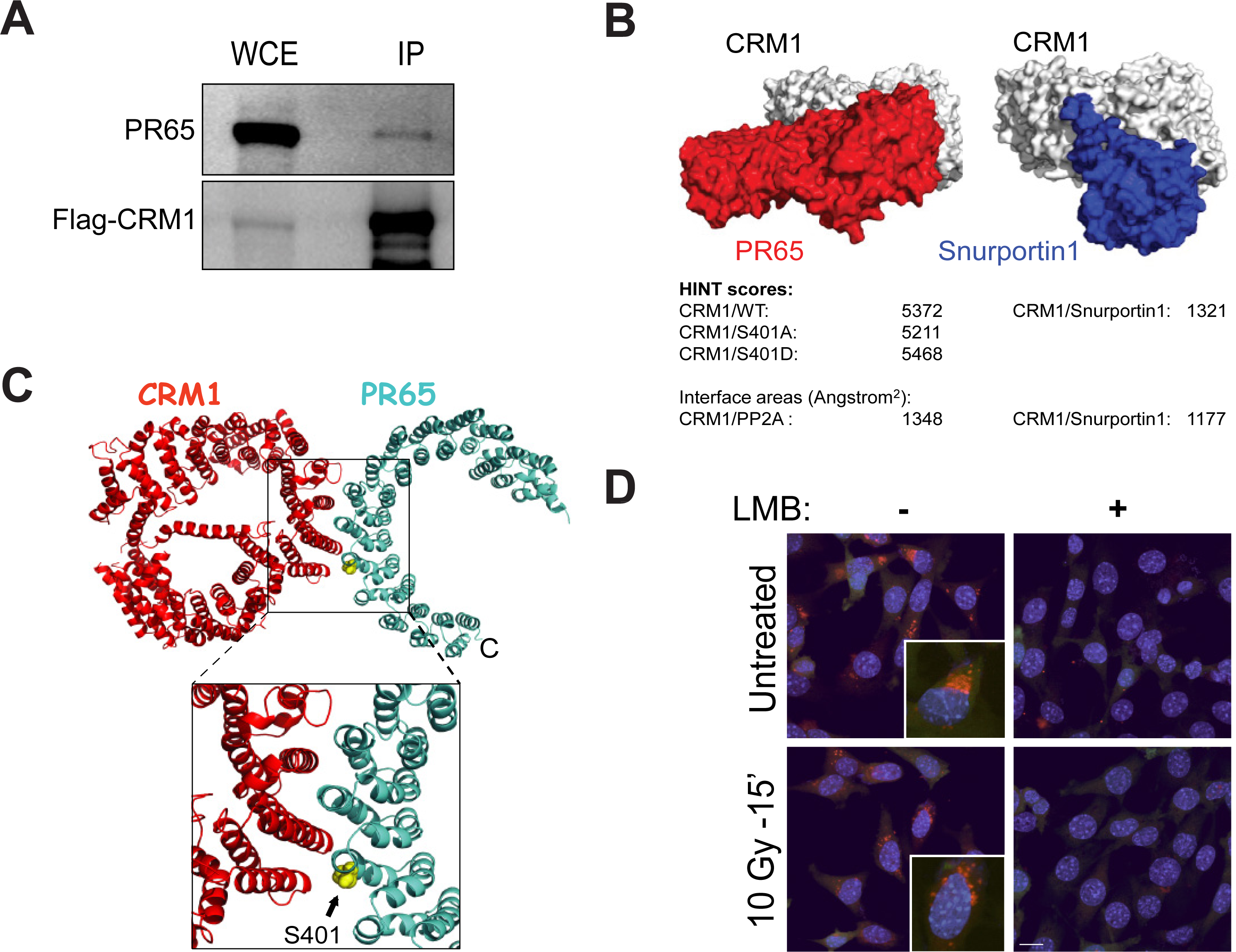
PR65 and CRM1 physically interact. (**A**) CRM1 forms a complex with PR65 as shown by co-immunoprecipitation with Flag-CRM1. HEK293 cells were transfected with Flag-CRM1, mRuby2-PR65 and RanQ69L expression plasmids followed by whole-cell immunoprecipitation at 72 hours using Flag-beads and western blotting with anti-PR65 and -Flag antibodies. (**B**) HINT analysis of CRM1-PR65 and CRM1-Snurportin 1 interactions suggest that CRM1-PR65 hydrophobic interaction is stronger than between CRM1 and Snurportin 1. (**C**) Modeling suggests that CRM1 (red) interacts with the - L_365_LPLFLAQL_373_-motif of PR65 (cyan) with S401 in the vicinity of the binding interface (yellow spheres). (**D**) PR65 and CRM1 co-localize at the cytoplasmic-nuclear membrane interface. Proximity Ligation Assay (PLA) shows that PR65 and CRM1 interact within 30-40 nm prior to irradiation with 10 Gy. After irradiation, this interaction was more punctated at the cytoplasmic-nuclear membrane. Pre-treatment with LMB (1 ng/ml) eliminates any sign of PR65-CRM1 interaction Scale bar is 10 μm.

We found that the docked CRM1-PR65 model has a HINT score more than 4-fold higher than CRM1-Snurportin-1 that is reduced when WT is replaced with S401A and increased with S401D. In addition, the CRM1-PR65 interface area was significantly larger compared with CRM1-Snurportin-1 (**Fig. 8B**). Furthermore, modeling suggests an interaction between CRM1-K537 and the -L_365_LPLFLAQL_373_-motif of PR65 with S401 positioned in the vicinity of the binding interface (**Fig. 8C**). Upon S401 phosphorylation, a tighter interaction is expected between K537 and pS401, or alternatively S401D, because of increased electrostatic interaction (**Supplemental Fig. 6B**). S401 is close to the surface of PR65 in the PP2A holoenzyme and at the interface where PR65 interacts with the PP2A catalytic (C) subunit between HEAT repeat 10-15 (**Supplemental Fig. 6C**). Interestingly, the putative PR65 NES is located in the inter-repeat loop between HEAT domains 10 and 11 and in close proximity to S401 within HEAT repeat 10 (106), and close to the outside boundary of the PP2A holoenzyme possibly more accessible for phosphorylation by ATM kinase (**Supplemental Fig. 6C**).

To confirm that PR65 and CRM1 interact in the cell, we carried out a proximity ligation assay (PLA). PR65-CRM1 proximity was observed as a fluorescent signal in untreated and irradiated WT cells with more punctate perinuclear appearance after irradiation. As expected, the PR65-CRM1 interaction was disrupted when cells were pretreated with LMB (**Fig. 8D**). Altogether, these results suggest that PR65 is exported to the cytoplasm by CRM1 after S401 is phosphorylated.

Because PP2A is such an abundant protein constituting *∼*1% of total protein in a mammalian cell and involved with regulating many processes throughout the cell it is technically challenging to follow a PR65 sub-population engaged in one specific process out of many in the cell using immunostaining or standard fluorescence protein fusions because of preexisting background. Therefore, we transfected photoactivatable plasmid constructs pPAmCherry-PR65 WT, S401A and S401D into HEK293 cells and demonstrate similar expression levels as endogenous PR65 (**Supplemental Fig. 7**). Laser photo-activation to a specific sub-cellular location such as the nucleus would allow for following the shuttling of PR65 in live cells after DNA damage with less interference from unrelated PR65 events. Forty-eight hours post transfection a sub-nuclear Region-of-Interest (ROI) was photo-activated with a 405 nm laser to activate PAmCherry-PR65 WT. We determined that laser alone was sufficient to cause DNA damage without including Hoechst 33528 to the medium based on γ-H2AX and phospho-KAP1 (S824) foci formation although the addition of dye increased DNA damage and seemed to enhance shuttling (**Fig. 9A**, **Supplemental Fig. 8**) with pan-nuclear PAmCherry-PR65 WT migrating to the nuclear membrane within minutes after photo-activation (**Fig. 9B, C**). We then confirmed our previous results with Flag-immunofluorescence (see **Fig. 7**), now using PAmCherry as reporter, that PR65-WT was exported from the nucleus to the cytoplasm whereas S401A was not, and PR65-S401D remained in the cytoplasm (**Fig. 9D, E, Supplemental** **Fig. 9**). We plotted the nuclear to cytoplasmic ratios (N/C) of the collected fluorescence signals. With WT cells, the N/C was as high as 0.05 initially and declined over several minutes (**Fig. 9E**, **Supplemental Table 2**), suggesting that PAmCherry-PR65 was nuclear early after DNA damage and photo-activation and quickly shuttled to the cytoplasm. The N/C ratio in the S401A and the S401D cells were initially 30-60% less (0.015-0.03) and remained the same throughout the experiment, suggesting that PR65-S401A and -S401D were statically nuclear and cytoplasmic, respectively (**Fig. 9E**, left panel). These results are consistent with the notion that nuclear PR65 is exported to the cytoplasm after DNA damage in a pS401-dependent manner. Additionally, in support of the physical presence of mutant PR65 in nucleus or cytoplasm, presumably with altered PP2A substrate specificity, is the distinct pattern of survival of MEFs after transfection with plasmids overexpressing different B regulatory subunits (**Supplemental Fig. 10**). We found a similar pattern of tolerability when B55α, B56α, or B56β were overexpressed in WT and S401 mutant cells, whereas a dramatically varied response was seen with B56γ1, B56γ3, B56*δ*, and B56*ε*, suggesting that PR65 has preferential B subunit partners depending on S401 status.

**Figure 9.**
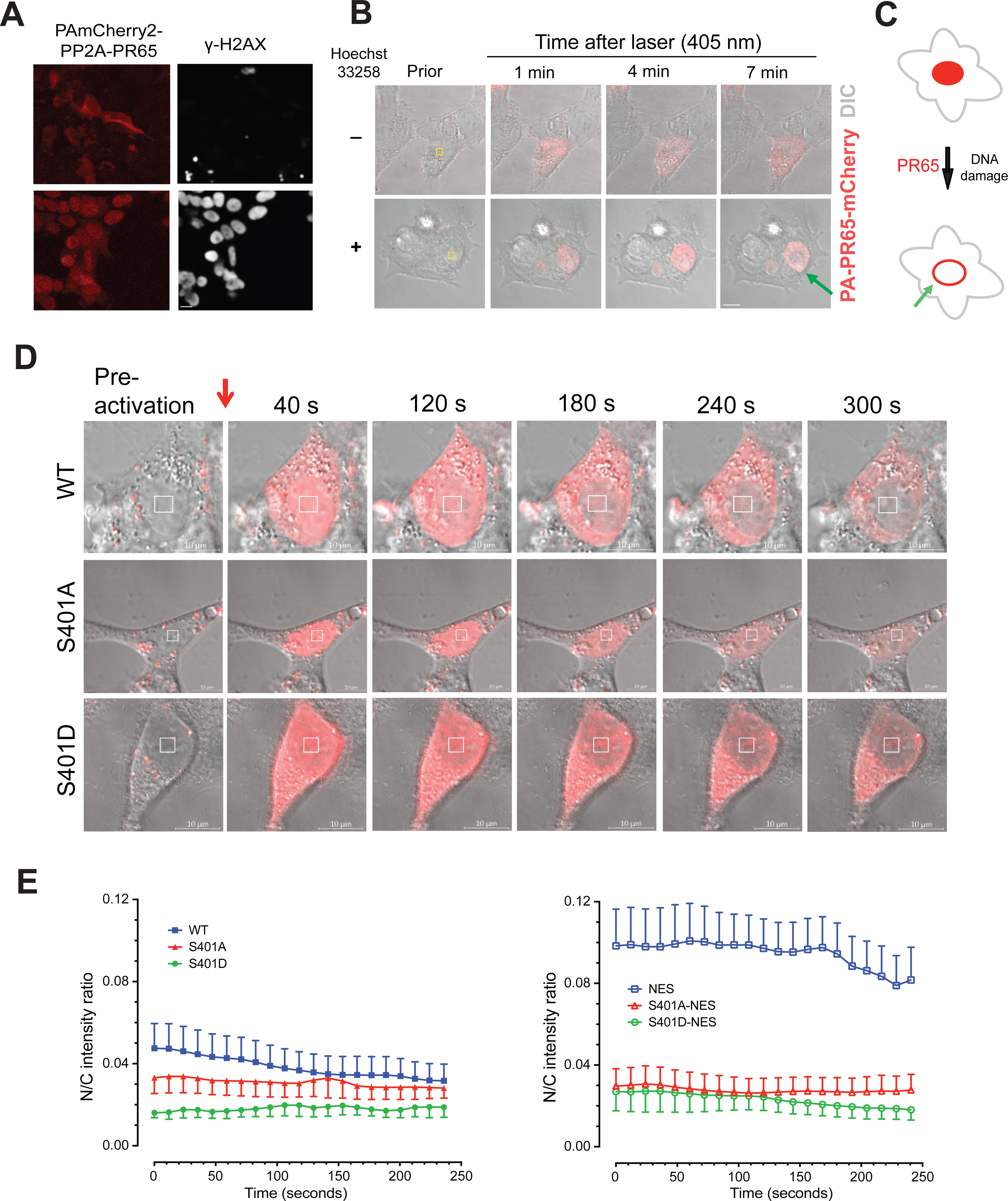
DNA damage triggers PR65 movement to the nuclear membrane. (**A**) Laser photo-activation of nuclear PR65 in HEK293 cells - adding Hoechst 33258 results in increased DNA damage and DSBs as indicated by the increased γ-H2AX nuclear staining Scale bar is 10 μm. (**B**) Nuclear PR65 migrates to the nuclear membrane after DNA damage. Time-lapse video shows that PR65 within minutes after 405 nm laser sub-nuclear irradiation (white square) is; 1) activated (mCherry signal increases) and 2) migrates to the nuclear membrane. The inclusion of Hoechst 33258 seems to increase the speed by which this movement occurs (green arrow) Scale bar is 10 μm. (**C**) Simplified model for PR65 accumulation at the nuclear membrane and export via the CRM1 nuclear protein complex. (**D**) pPAmCherry-PR65-WT is exported to the cytoplasm after nuclear irradiation. HEK293 cells were transfected with pPAmCherry-PR65 (WT, S401A and S401D) in glass bottom dishes and 48 h post-transfection sub-nuclear ROI (4 μm x 4 μm), marked by the squares, were photoactivated by 405 nm laser (242.8 μJ) and followed over time (< 300 s) by live cell imaging using a Zeiss LSM 710 microscope with images acquired at 63x. WT-PR65 is exported from nucleus to cytoplasm over time whereas PR65-S401A is retained in the nucleus and PR65-S401D remains cytoplasmic Scale bar is 10 μm. (**E**) PR65 NES mutation dramatically alters the behavior of PR65. In WT cells, the N/C was as high as 0.05 initially and declined over several minutes, suggesting that PAmCherry-PR65 was nuclear early after DNA damage and photo-activation and quickly shuttled to the cytoplasm. The N/C ratio in the S401A and the S401D cells were initially 30-60% less (0.015-0.03) and remained the same throughout the experiment, suggesting that PR65-S401A and -S401D were statically nuclear and cytoplasmic, respectively (left panel). When HEK293 cells were transfected with pPAmCherry-PR65 WT/L373A and photo-activated, the initial N/C fluorescence intensity ratio was 0.10, twice the ratio seen with WT, and the signal did not decline over time except at >170 s that might represent photo bleaching. The S401A and S401D mutants were less affected by the L373A addition, but, interestingly, the flat N/C ratios seem to come more together compared with S401 alteration alone (right panel). One-way ANOVA was performed on the time series. p<0.0001 for WT vs S401A, WT vs S401D, NES vs NES-S401A and NES vs NEs-S401D.

**Figure 10.**
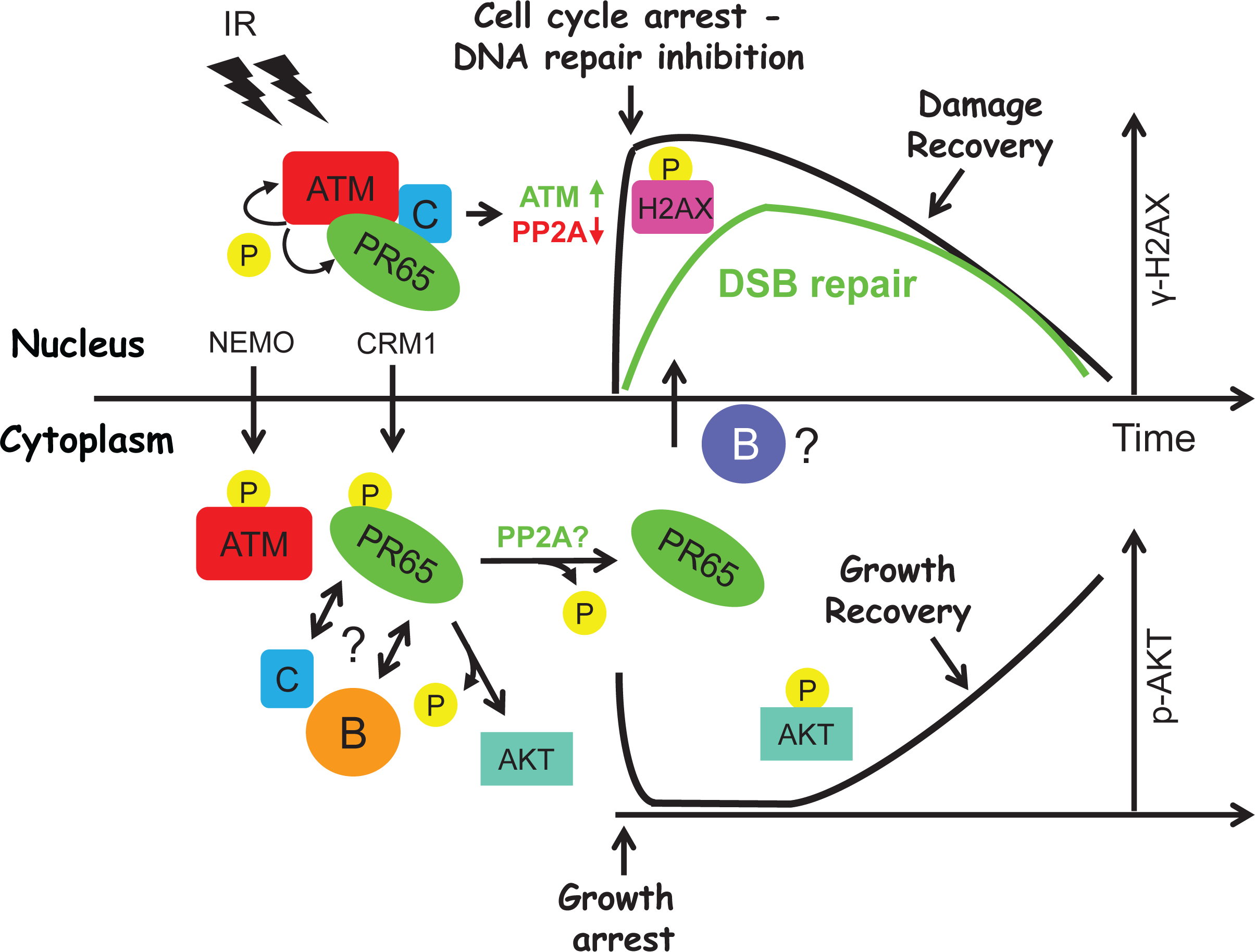
Model for ATM-PP2A mediated regulation of the DDR and cell growth - spatio-temporal (de)phosphorylation and nuclear-cytoplasmic shuttling by ATM and PP2A controls the DDR “ON-OFF” switch. ATM phosphorylates many proteins in response to IR including H2AX at S139 and PR65 at S401 thereby initiating the DDR and triggering cell cycle checkpoints and growth arrest, and transiently inhibiting DSB repair. Within minutes, phosphorylated PR65 translocates to the cytoplasm that reduces PR65 activity in the nucleus allowing the DDR with its highly phosphorylated protein landscape to run its course. At the same time, AKT and other growth-promoting factors and signaling pathways are inhibited by the increased PR65 activity in the cytoplasm while the damage is assessed and repaired. Soon after the DDR is beginning to subside, PR65 might shuttle back to the nucleus where it increases PR65 activity and begin to engage DSB repair while γ-H2AX and other DDR targets are dephosphorylated. When repair is completed and cells resume cycling, growth recovers.

To determine whether the putative PR65 NES sequence functions, we generated photoactivatable plasmid constructs, pPAmCherry-PR65 WT, S401A, and S401D, also having the L373A alteration. When HEK293 cells were transfected with pPAmCherry-PR65 WT/L373A and then photo-activated, the initial N/C fluorescence intensity ratio was 0.10, twice the ratio seen with WT, and the signal did not decline over time except at >170 s that might represent photo bleaching (**Fig. 9E**, right panel, **Supplemental Table 2**). The S401A and S401D mutants were less affected by the L373A addition, but, interestingly, the flat N/C ratios seem to come more together compared with either S401 alteration alone. Altogether, our results show that ATM kinase serves as a gatekeeper for regulating the initial DDR by phosphorylating PR65 at S401 exposing a buried NES to CRM1 and resulting in nuclear export of PR65.

## DISCUSSION

PP2A plays a major role in numerous cellular processes and has been associated with regulating the DDR and DNA repair (25,27,29,30). ATM kinase phosphorylates PR65 at S401, which is associated with dysfunctional neuronal chromatin deacetylation by HDAC4 resulting in neurodegeneration in ataxia telangiectasia (31). Previous work by our group has explored how DSB signaling through ATM converges on pro-survival signaling through AKT (21). Negative regulation of AKT by PP2A is well-established (107, 108). Herein we report that phosphorylation of a single, critical amino acid residue (S401) in the PR65 subunit is important for cell survival, DNA damage signaling, and DNA repair.

Our molecular modeling simulations suggest that phosphorylation of S401 causes the horseshoe-shaped scaffolding subunit PR65 to undergo a conformational change. This leads to the disassociation of the ATM-PR65 complex, as previously reported (25), which, herein is supported by our co-immunoprecipitation results. We also show that ATM, AKT and PP2A-C no longer interact with S401-phosphorylated PR65 or the analogous phospho-mimetic mutant. Impaired interaction of S401D with PP2A-C could therefore fundamentally alter PP2A phosphatase activity, substrate targeting, and/or cellular localization. A reciprocal immunoprecipitation experiment might resolve whether AKT is part of the PR65-ATM complex or is a separate PR65-AKT complex.

Interestingly, PR65 and ATM both have many HEAT repeats, 15 and 49, respectively. HEAT repeats consist of pairs of interacting, anti-parallel helices linked by flexible “intra-unit” loops believed to form dynamic structures and elastic connectors that link force and catalysis (106, 109). Additionally, CRM1 has 19 HEAT repeats known to participate in the formation of a nuclear export complex (110). It is tempting to speculate that these HEAT repeats are involved in a dynamic “hand-over and exchange” process swapping partners during the DDR and resulting in nuclear export of PR65 expected to quickly alter the nuclear phospho-protein landscape and maintain a state of elevated phosphorylation in the nucleus immediately following DNA damage. Presumably, PR65 then shuttles back into the nucleus to turn off the DDR by dephosphorylating proteins targeted directly or indirectly by ATM phosphorylation. Clearly, this is a simplified scenario since several other S/T protein phosphatases such as PP1, PP4 and PP6 have also been linked to the DDR (111). However, it is well established that these phosphatases perform redundant functions by acting either as the primary phosphatase or secondary backup to avoid cellular crises to occur (e.g., during mitosis).

We, and others, have previously reported that an ATM-dependent phosphatase negatively controls AKT (S473) dephosphorylation in response to insulin and radiation (21, 112). We hypothesized that ATM mediated phosphorylation of PR65 at S401 might alter PP2A phosphatase activity thereby increasing AKT S473 phosphorylation after such stimulation which can be blocked by an ATM kinase inhibitor (21). In the present study, we found that AKT phosphorylation was elevated in S401D mutants relative to the WT and S401A counterparts. However, in response to radiation, the increase in AKT (S473) phosphorylation was impaired in S401D as well as S401A cells. We know that the DDR initially takes place primarily in the nucleus, whereas insulin signaling initiates at the plasma membrane through activation of insulin receptor and with AKT present in both compartments. It is likely that AKT is differentially regulated in the nucleus and cytoplasm by PP2A. This could be due to reduced PP2A activity and/or inappropriate cellular location of PP2A impairing the normal execution of the DDR. That the response to insulin and radiation are different might be expected because radiation likely produce more pleiotropic effects on signaling than insulin (69). This may be through association with different B subunits, and as demonstrated herein, in a coordinated PP2A nuclear-cytoplasmic shuttling process to spatio-temporally regulate the DDR as well as cell growth.

γ-H2AX is present at DSBs in the nucleus. From our PR65 localization study, we observed that S401D was retained in the cytoplasm in cells exposed to radiation whereas the WT, and to a lesser extent S401A, was found in both the cytoplasm and nucleus. Thus, cytoplasmic sequestration of PR65-S401D possibly makes cells impaired in nuclear γ-H2AX dephosphorylation. We have also shown that S401D cells are less efficient in DSB repair, accumulate chromosomal abnormalities and show reduced survival after DNA damage. On the other hand, S401A cells have much greater end joining activity compared to WT cells *in vitro,* which is in line with the idea that DNA repair cannot shut off. We speculate that this is due to the inhibition of nuclear export of PR65. Furthermore, since PP2A is essential for the radiation G2/M checkpoint (29), the decreased ability of PR65 to shuttle and retain proper PP2A activity in a strict spatiotemporal manner would likely result in checkpoint failure and mitotic catastrophe.

Most notably, both *in vitro* and *in vivo* studies suggest that DSB repair is fundamentally altered in the S401 mutants. Future studies will determine in more detail what type of DSB repair is engaged when both c-NHEJ and HRR are compromised as is the case in S401D cells. Most likely, a micro-homology type of DSB repair such as TMEJ or SSA is involved to rescue cell survival based on our preliminary analyses.

As to the mechanism how PR65 S401 alterations might affect cell growth and DNA repair in a reciprocal manner, we propose a model in which PP2A together with ATM might serve as gatekeeper of the DDR (**Fig. 10**). For ATM-directed activation of the DDR to occur through a phosphorylation cascade, the phosphatase, in this case PP2A, must initially be inactivated to maintain a phosphorylated protein landscape that subsequently has to be reactivated by removing phosphates to reenter the cell cycle and resume growth. If the DNA damage is sub-lethal, DNA repair is mostly completed within a relatively short time window of 6-12 hours (113). Interestingly, ATM and PP2A form an auto-regulatory, yin yang/kinase-phosphatase relationship serving as a potential on-off switch of the DDR. Based on our findings and by others, PP2A is likely having the upper hand in this relationship since inhibitors of PP2A activate the ATM kinase, resulting in a pseudo-DDR response without significant DNA damage (25). There are many examples of this regulatory hierarchy in kinase-phosphatase signaling pathways (114). Nuclear export seems a fast, efficient means of temporarily removing PP2A from sites of DNA repair. Once repair is complete, PP2A can shuttle back to the nucleus to remove protein phosphates and reset the cell for baseline growth and activity. Therefore, PP2A (along with other phosphatases) must coordinate and balance DNA repair and cell survival with apoptosis resulting from insurmountable cellular damage. Our model incorporates the ATM-PP2A “on-off” switch for the DDR coupled with cell recovery and growth by ATM and PP2A shuttling between the nucleus and cytoplasm and, together, acting as a cellular rheostat for DNA damage and growth control.

Our results suggest that the improper localization of PP2A/PR65 and, thus, untimely dephosphorylation of critical phospho-proteins result in a shift in the quality of DSB repair though without dramatic impact on survival. The most straightforward interpretation of our results is that DSB repair is re-wired in a manner that substitute c-NHEJ and HRR with some other homology-based DNA repair to maintain viability after DNA damage. Regardless, because the only difference between the three cell populations used in our study is the PR65 S401 alterations, this cell system could be a rich source for dissecting DNA repair and growth responses regulated by PP2A that would otherwise not be possible with general DNA damaging treatments, which would trigger a plethora of difficult-to-dissect cellular responses.

## Acknowledgements

Services in support of the research project were provided by the VCU Massey Cancer Center Transgenic/Knockout Mouse Core, the Microscopy Shared Resource, the Flow Cytometry Shared Resource, and the Cancer Mouse Model Core supported, in part, with funding from NIH-NCI Cancer Center Support Grant P30 CA016059.

## Reporting summary

Further information on research design is available in the Nature Research Reporting Summary linked to this article.

## Data availability

All the data supporting the findings of this study are either available within the paper and its Supplementary Information files or can be obtained from the authors upon request.

## Supplemental data

### Materials and Methods

#### Plasmids

Plasmids:
| Plasmids | Source | Provided by |
| --- | --- | --- |
| pMIG-Aalpha | Addgene #10884 | William Hahn |
| Flag-hCRM1 | Addgene #17647 | Xin Wang |
| pmCherry-C1-RanQ69L | Addgene #30309 | Jay Brenman |
| pCSCMV:tdTomato | Addgene #30530 | Gerhart Ryffel |
| Human TERT (LOX-TERT-iresTK | Addgene #12245 | Didier Trono |
| pcDNA3-mRuby2 | Addgene #40260 | Michael Lin |
| pGEX-2T | GE Life Sciences #28954653 |  |
| YFP-ATM |  | David Chen |
| pBabe-hygroPyMT | Addgene #22305 | Filippo Giancotti |
| pPAmCherry-a-tubulin | Addgene #31930 | Vladislav Verkhusha |
| LentiCRISPR v2 | Addgene plasmid # 52961 | Feng Zhang |
| LentiCRISPRv2 hygro | Addgene plasmid # 98291 | Brett Stringer |
| pCEP-4HA-B56 alpha | Addgene #14532 | David Virshup |
| pCEP-4HA-B56 gamma3 | Addgene #14535 | David Virshup |
| pCEP-4HA-B56 gamma1 | Addgene #14534 | David Virshup |
| pCEP-4HA-B56 epsilon | Addgene #14537 | David Virshup |
| pCEP-4HA-B56 delta | Addgene #14536 | David Virshup |
| pCEP-4HA-B56 beta | Addgene #14533 | David Virshup |
| pCEP-4HA-B55 alpha | Created by replacing B56 alpha from pCEP-4HA-B56 beta with B55 alpha. The B55 alpha cDNA was obtained from pUC19-hPPP2R2A by PCR. | This study |

#### Antibodies and Reagents

**Antibodies and Reagents:**
| **Antibodies and Reagents** | **Company** | **Catalog no.** |
| --- | --- | --- |
| anti-ATM | Cell Signaling Technology | 2873 |
| anti-AKT | Cell Signaling Technology | 2920 |
| anti-p(S473) –AKT | Cell Signaling Technology | 9271 |
| anti-PP2A-C | Cell Signaling Technology | 2038 |
| anti-pS1390-gH2AX | Cell Signaling Technology | 9718 |
| anti-GST | Cell Signaling Technology | 2624 |
| anti-γ-H2AX | Millipore Sigma | 05-636 |
| anti-GAPDH | Millipore Sigma | MAB374 |
| anti-Flag | Millipore Sigma | F1365 |
| anti-53BP1 | Novus Biologicals | NP100-304 |
| anti-Rad51 | Novus Biologicals | NB100-148 |
| anti-PyMT | Novus Biologicals | NB100274955 |
| anti- p(S4/S8)-RPA | Bethyl laboratories | A300-245A |
| anti-ERK2 | Santa Cruz Biotechnology | sc-154 |
| anti-pERK1/2 | Santa Cruz Biotechnology | sc-7383 |
| anti-CRM1 | Santa Cruz Biotechnology | sc-74454 |
| Alexa Fluor 594 goat anti-mouseIgG | Life Technologies | R37121 |
| Alexa Fluor 680 goat anti-rabbit IgG | Life Technologies | A-21076 |
| Alexa Fluor 680 goat anti-mouse IgG | Life Technologies | A-21057 |
| Dylight 800 anti-mouse IgG | Cell Signaling Technology | 5257 |
| Dylight 800 anti-rabbit IgG | Cell Signaling Technology | 5151 |
| Leptomycin B | Millipore Sigma | L2913 |
| Camptothecin | Millipore Sigma | 7689-03-4 |
| KU-60019 | Selleckchem | S1570 |

#### Immunoprecipitation

Flag-PR65 was immunoprecipitated from HEK293 cells stably expressing pMIG-Aalpha WT, S401A or S401D with anti-Flag M2 beads (Sigma Aldrich, Cat #F2426) in lysis buffer with Halt™ protease and phosphatase inhibitor cocktail (ThermoFisher Scientific, Cat #78440). YFP-ATM was immunoprecipitated by using the GFP-Trap Magnetic Agarose beads (Chromotek, Cat #gtma-10).

#### Photoactivation assay image processing

Temporal averages of the series were used for initial segmentation of cytoplasmic regions where mCherry fluorescence was stronger than in the corresponding irradiated nuclei. Segmentation thresholds were calculated by adding average and standard deviation (multiplied by 0.3) of fluorescence in the irradiated nuclear regions. The resulting binary masks were inverted and used to limit conditional dilation of the seeds (10x10 pixels) corresponding to the irradiated regions. Next the initial nuclear masks, created in this manner, were used construct 3-pixel wide regions around them, at the distance of 4 pixels. Products of these regions and the result of the initial cytoplasm segmentation were used as cytoplasmic counterparts of the nuclear masks. These initial masks were updated tor every image in the post-irradiation series by conditional binary closing (2 iterations) of nuclear masks and according to construction of the cytoplasmic masks. Median intensity of the pre-irradiation images was used as background estimator. Average nuclear and cytoplasmic mCherry fluorescence intensities were corrected for background. The respective nucleus/cytoplasm ratios were averaged over cell populations.

#### Proximity Ligation assay

Proximity ligation assay was performed as per manufacture’s recommendations (Duolink^®^ using PLA^®^ technology from Sigma Aldrich). PR65 KO MEFs expressing Flag-tagged WT PR65 were seeded on Lab-Tek (Naperville, IL) glass slides and treated with Leptomycin B (2 ng/ml) or not for 3 hours, irradiated with 10 Gy or not, or left untreated. Cells were fixed after 15 min followed by rabbit anti-PR65 and mouse anti-CRM1 antibodies to determine the extent of the PR65-CRM1 interaction. Images were obtained by confocal microscopy.

### Supplemental Figure Legends

**Supplemental Figure 1.**
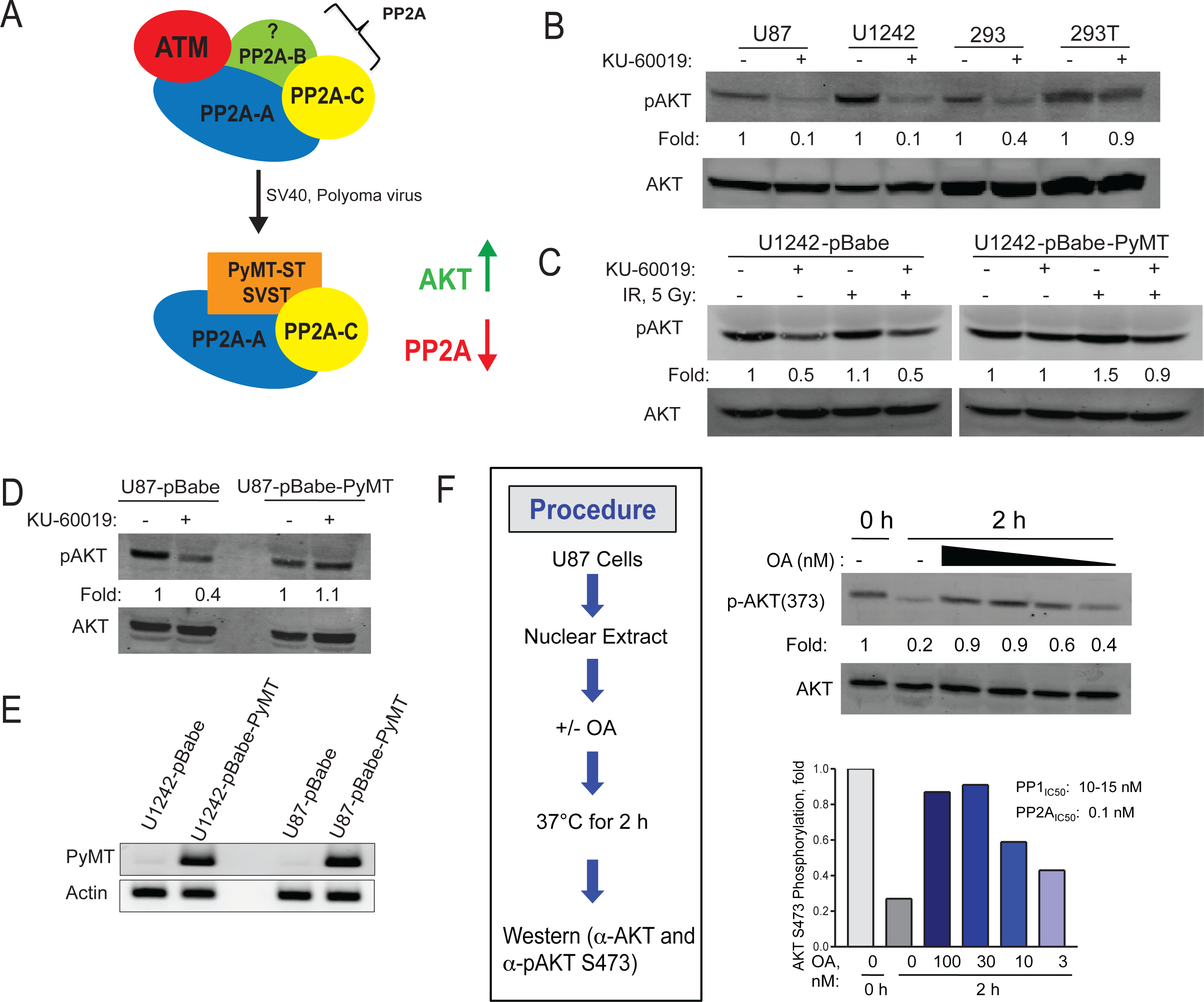
SV40 small-t and polyoma Middle-T inhibit PP2A and blocks radiation-induced AKT activation. **(A)** SV40 t (*SVST*) and Polyoma MT (*PyMT*) antigens are known to replace the PP2A-B subunit and bind to the PP2A-A and C Core enzyme to inhibit PP2A activity and increase AKT phosphorylation (1) **(B)** Inhibition of ATM does not reduce pAKT (S473) levels in HEK293T cells but does in U87, U1242, and HEK293 cells. U87, U1242, HEK293, and 293T cells were treated with or without KU-60019 (3 μM) for 1 hour. Whole cell extracts were separated on an SDS-PAGE gel and analyzed by western blotting using anti-pAKT (S473) and anti-AKT antibodies. **(C)** U1242 and **(D)** U87 cells expressing PyMT antigen show diminished ATM-mediated inhibition of pAKT (S473) levels after radiation. U1242 and U87 cells were infected with pBabe-hygroPyMT vector or control vector pBabe-hygro. Infected cells were pre-treated with or without KU-60019 (3 μM) for 1 hour followed by 5 Gy and whole cell extracts prepared at one hour post-IR were separated on an SDS-PAGE gel and analyzed by western blotting using anti-p-AKT and anti-AKT antibodies. **(E)** U1242 and U87 infected with PyMT virus are positive for virus integration. **(F)** Treatment of nuclear extract with okadaic acid increases pAKT levels. Nuclear extract was prepared from human glioma U87 cells and incubated with OA (3, 10, 30, or 100 nM) or not for 2 hours. Samples were then separated by SDS-PAGE and transferred to a membrane followed by sequential western blotting with anti-pAKT (S473) and AKT antibodies. Quantification of band intensity was done by densitometric scanning.

**Supplemental Figure 2.**
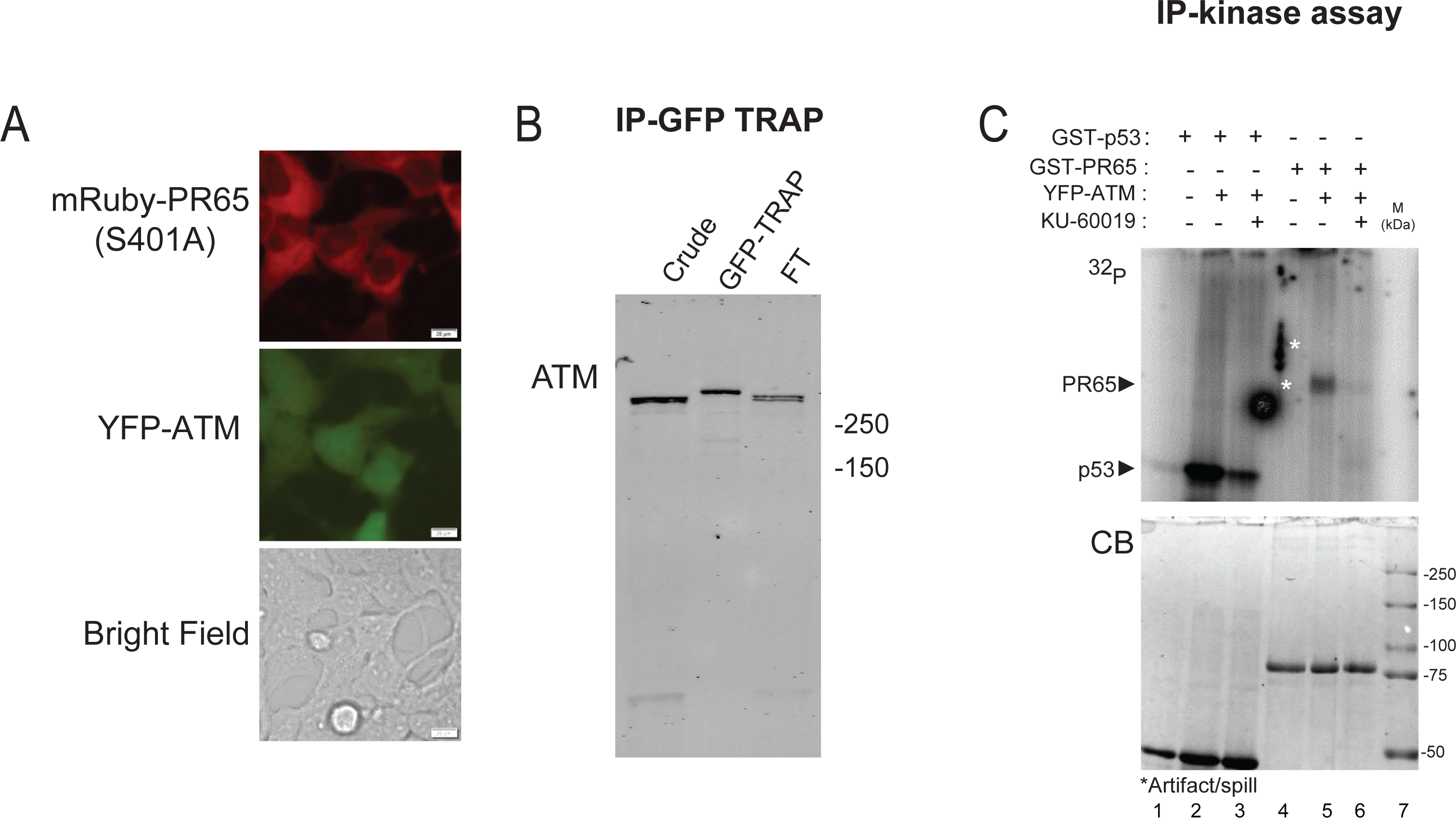
ATM kinase phosphorylates PR65 in vitro. **(A)** HEK293 cells were transfected with YFP-ATM +/- mRuby2-PR65 constructs and sorted with only YFP-ATM/PR65 (S401A) cells surviving suggesting that ATM and PR65 S401A, but not WT or D, can be stably co-expressed in HEK293 cells whereas S401 phosphorylation or expression of S401D phosphor-mimetic is not tolerated with over-expressed ATM. FT; flow-through. **(B)** Whole cell extracts of HEK293 cells expressing YFP-ATM were immunoprecipitated with GFP-TRAP (ChromoTek). **(C)** In vitro kinase assay was performed by suspending YFP-ATM immunoprecipitate w/o co-expressed PR65 in buffer with [γ-^32^P] ATP, 20 μM unlabeled ATP, GST-substrates (GST-PR65FL or GST-p53100) with or without KU-60019 (1 μM) for 1 hour. Samples were separated on a SDS-PAGE gel and then analyzed by a Typhoon phosphoimager (PerkinElmer) (*top panel*). Phosphorylation of p53 and PR65 is reduced in the presence of KU-60019 (lane 3 and lane 6) relative to controls (lane 2 and 5), respectively. Coomassie Blue (*CB*) - stained image of the gel showing GST-p53 and GST-PR65 proteins (*bottom panel*).

**Supplemental Figure 3.**
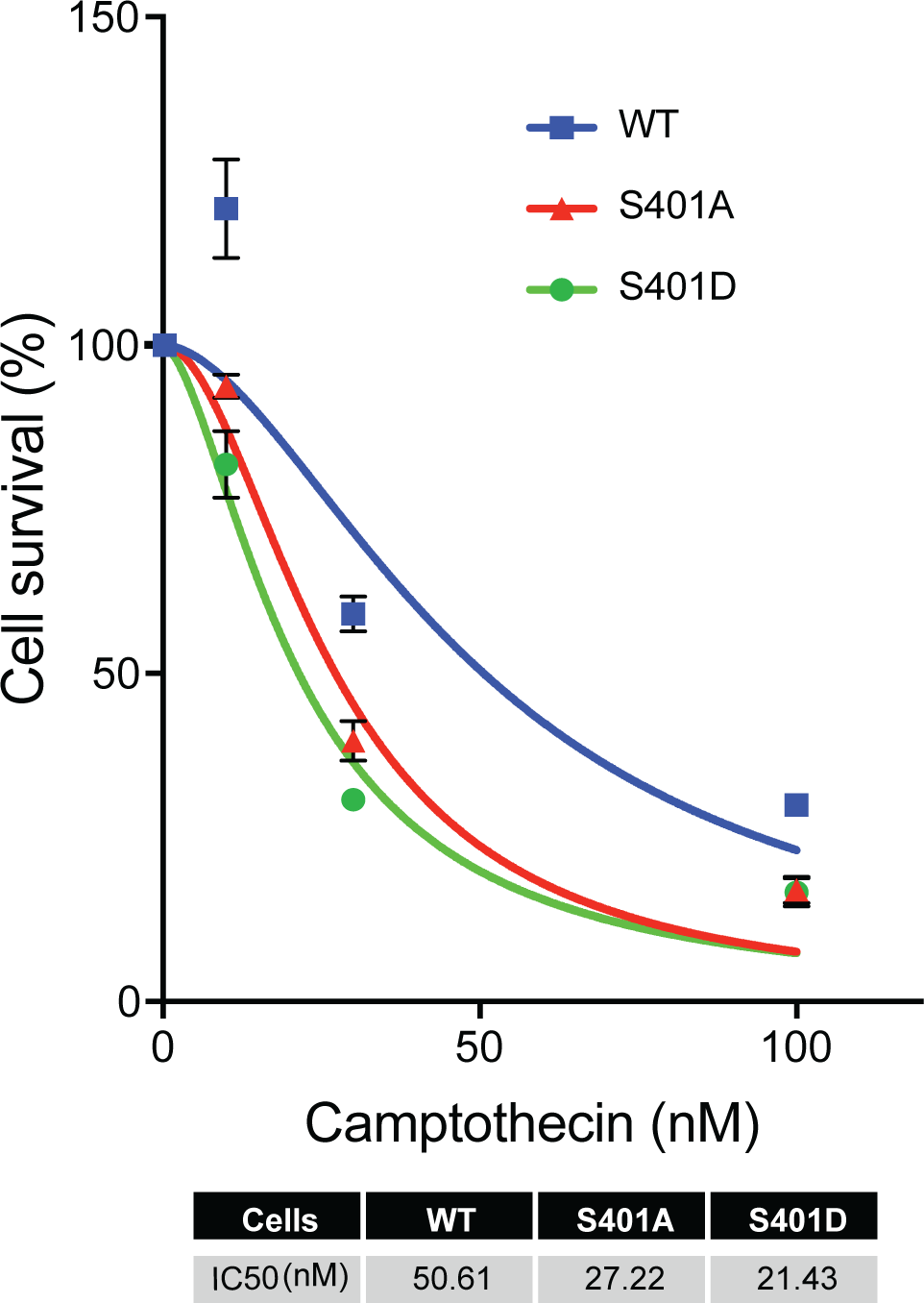
S401 mutant cells are more sensitive to camptothecin than wild type MEFs. WT, S401A and S401D MEFs were treated with CPT and analyzed by Cell titer-Glo® assay at 96 hours to determine cell death. Error bars; mean ± SEM (n = 3). CPT IC_50_ is presented in the table.

**Supplemental Figure 4.**
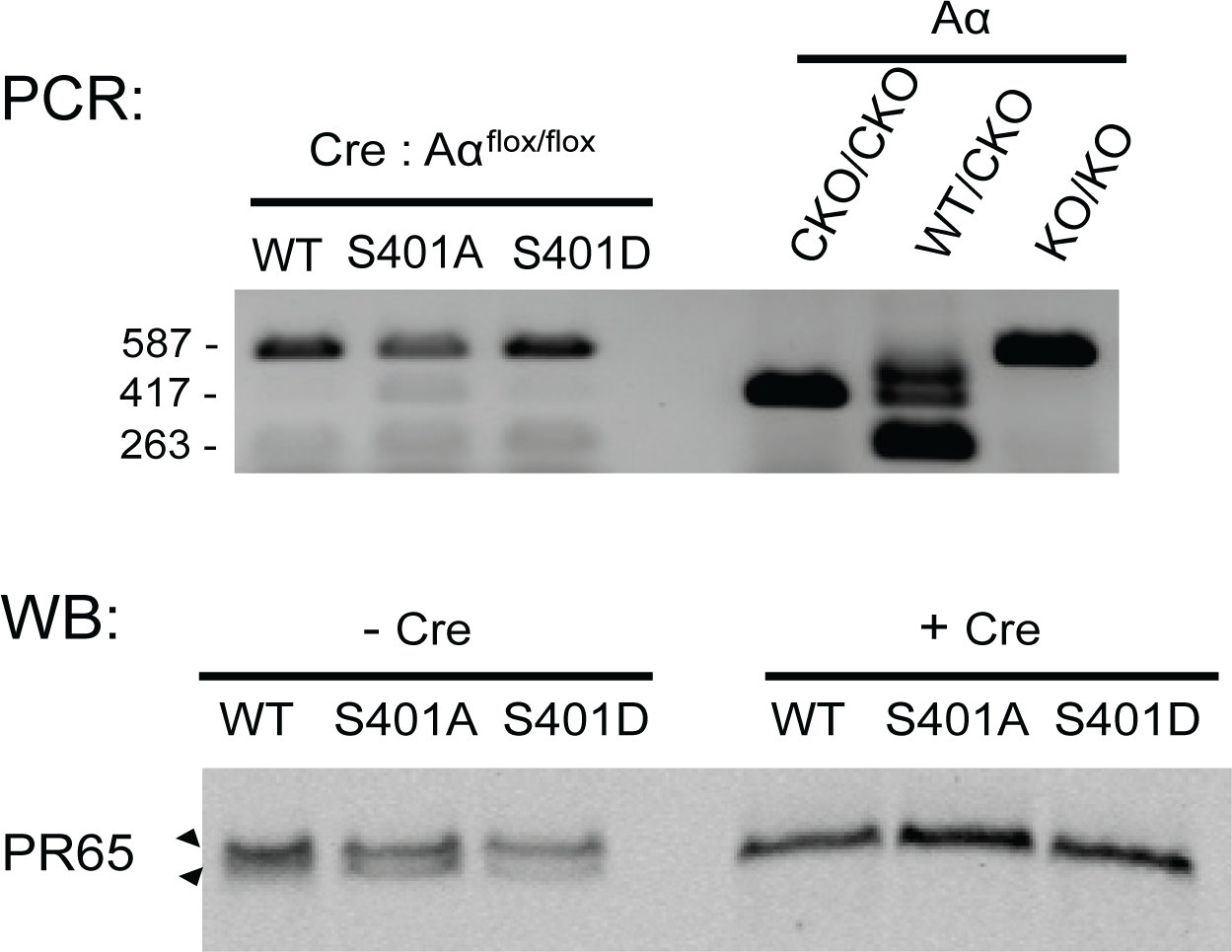
PCR and western blot analyses to verify floxing of endogenous PR65 sequences in MEFs stably transfected with DR-GFP. PCR screening with primers P1 and P3 generated a 263-bp product for the WT allele and a 417-bp product for the CKO allele. In cells where Cre was expressed, primer pair P3 and P2 generated a 587-bp product for the PR65 KO allele (*top panel*). Whole cell extracts of WT, S401A, and S401D MEFs with integrated DR-GFP infected with Ad-Cre or not were analyzed by western blotting with anti-PR65 antibody (*bottom panel*). Notice that the bottom PR65 band is missing after floxing indicating the loss of the endogenous allele. In addition, Flag-PR65 levels of WT, S401A, and S401 are very similar (compare with **Fig. 1E**).

**Supplemental Fig 5.**
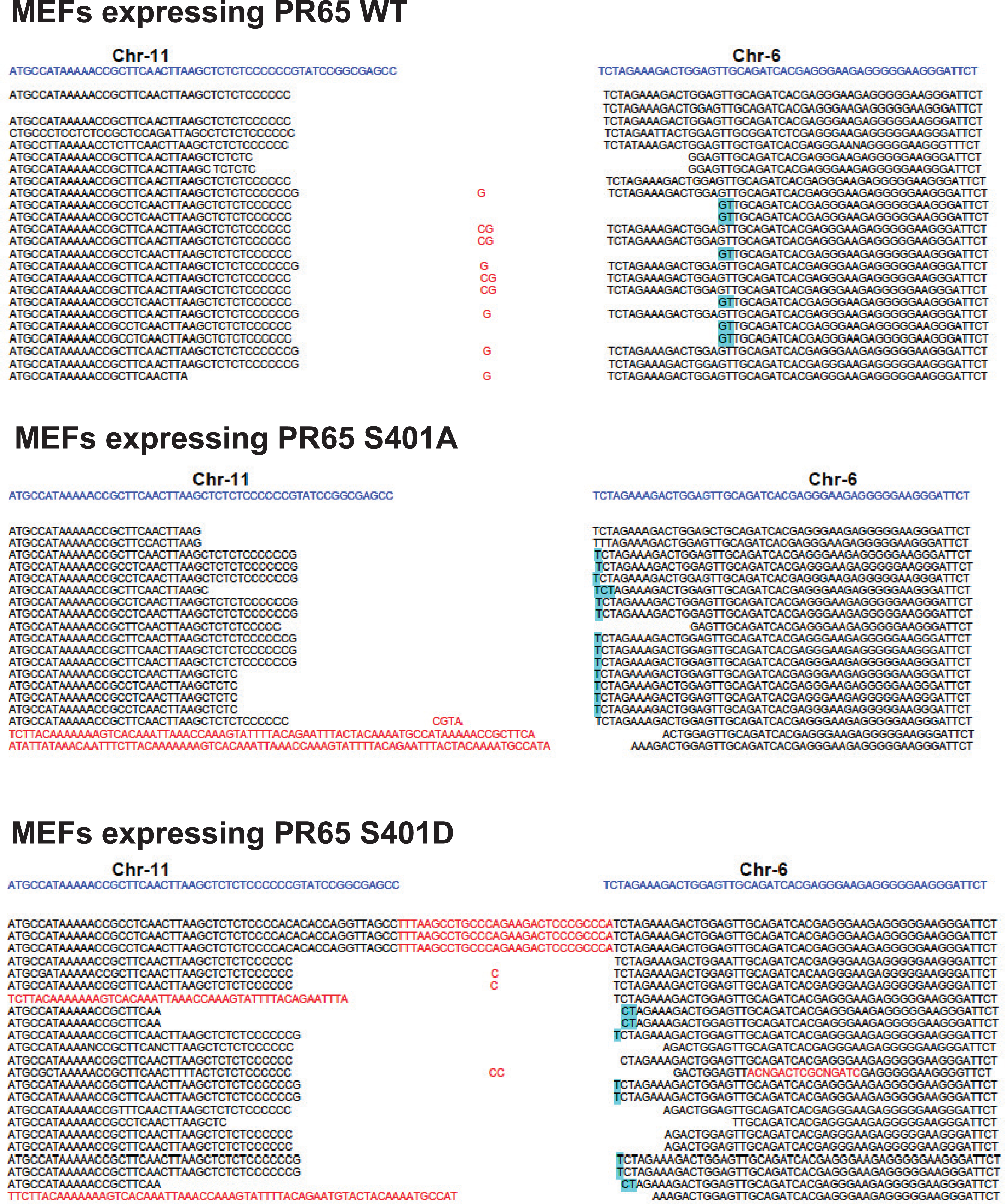
Translocation between DSBs (CRISPR-Cas9) at Rosa26 (Chr 6) and H3f3b (Chr 11). Sequences of Chr(11) breakpoint junction from WT **(A)**, S401A **(B)** and S401D **(C)** cells. Reference sequence is highlighted at the top (*blue*). The remaining DNA sequences represent individual translocations recovered by PCR and subject to Sanger sequencing. Nucleotide insertions are marked in red. Nucleotide deletions are represented as gaps. Micro-homology is denoted by cyan highlighted nucleotides.

**Supplemental Figure 6.**
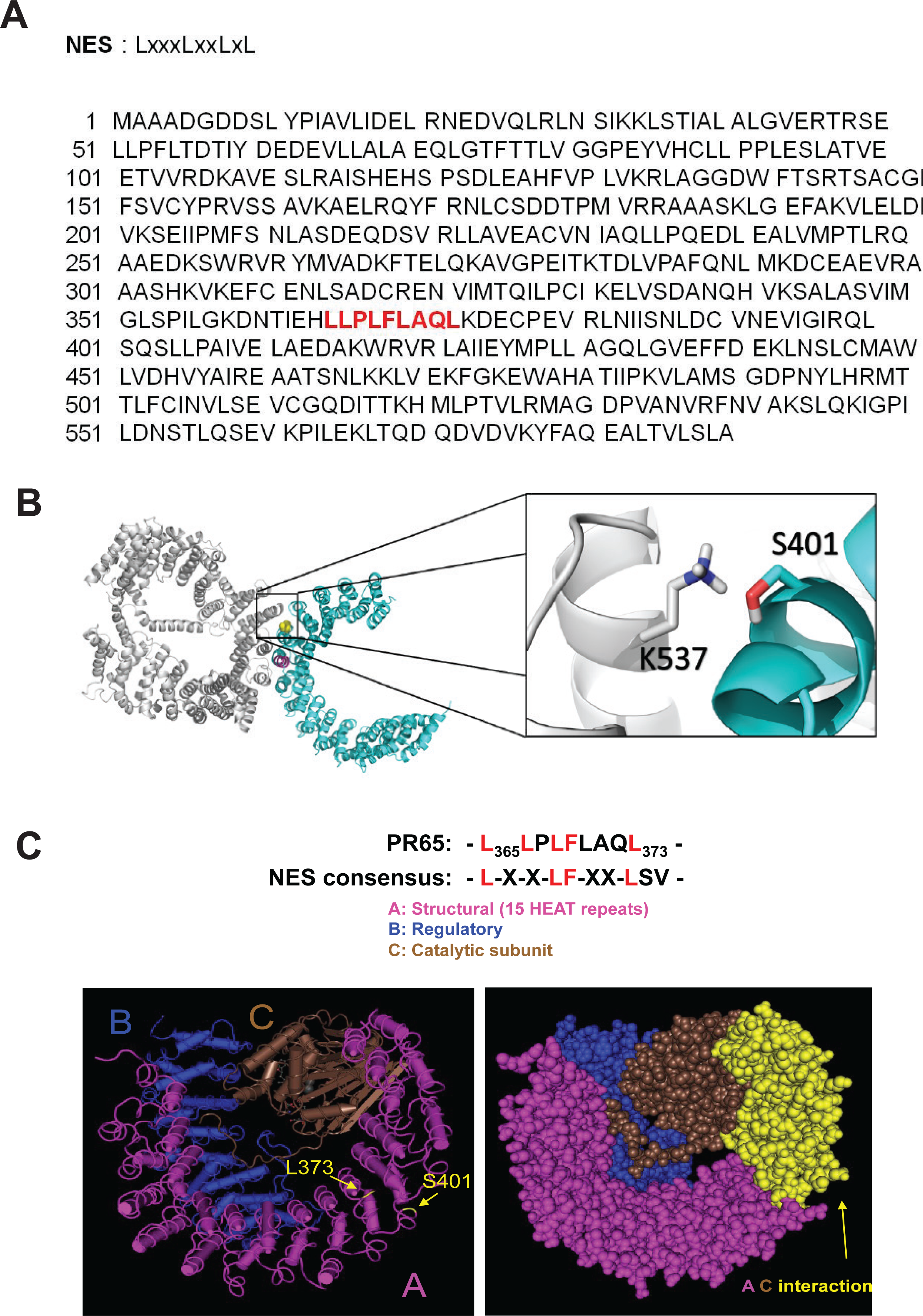
Putative PR65 NES in close proximity to S401. **(A)** PR65 has a putative nuclear export sequence (highlighted in red). **(B)** Model for PR65-CRM1 interaction via PR65-S401 and CRM1-K537. PR65-CRM1 co-crystal modeling showing a possible key interaction between PR65-S401 and CRM1-K537. PR65-CRM1 is colored in cyan whereas CRM1 is colored in silver. **(C)** PR65 S401 phosphorylation might expose a buried PR65 NES. A putative NES is located in the inter-repeat loop between HEAT domains 10-11 of PR65 and in close proximity to S401. Co-crystal structure of PP2A with PR65 structural (A), catalytic (C), and (B) regulatory subunits with S401 and putative NES at L373 shown in yellow (*left panel*). Interaction domains between PR65 (A) and catalytic (C) subunits (*right panel*). The *yellow* portion of PR65 marks the domain directly interacting with the PP2A catalytic subunit (*brown*). PR65-S401 is located at the rim whereas L373 is buried deeper at the interface between the PR65 and the PP2A catalytic subunit (*left panel*).

**Supplemental Figure 7.**
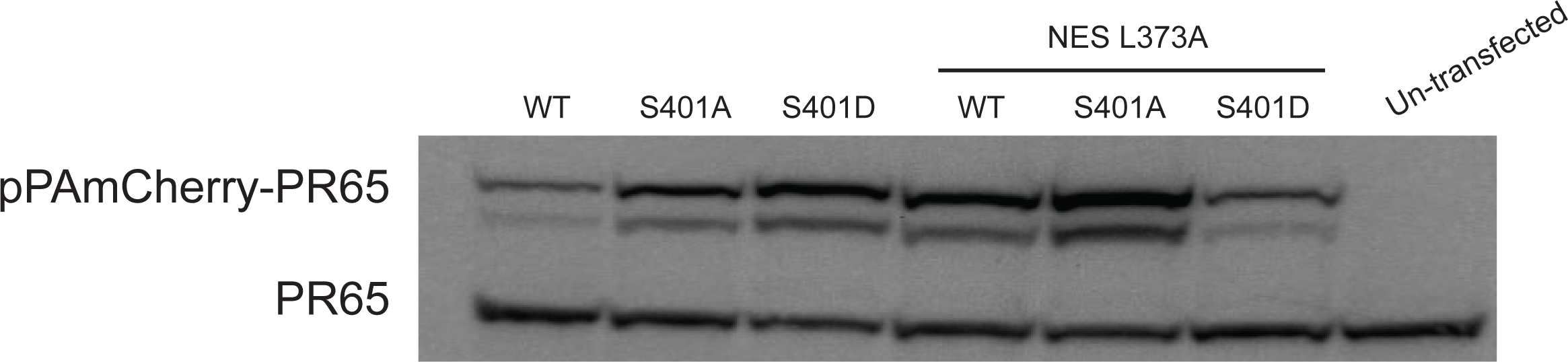
Expression of PR65-NES and S401A/NES and S401D/NES mutants. Transfection of HEK293 followed by western blotting of extracts shows approximately similar expression levels with anti-Flag and -PR65 antibodies. Notice the size difference between endogenous PR65 and much larger PAmCherry-PR65 proteins.

**Supplemental Figure 8.**
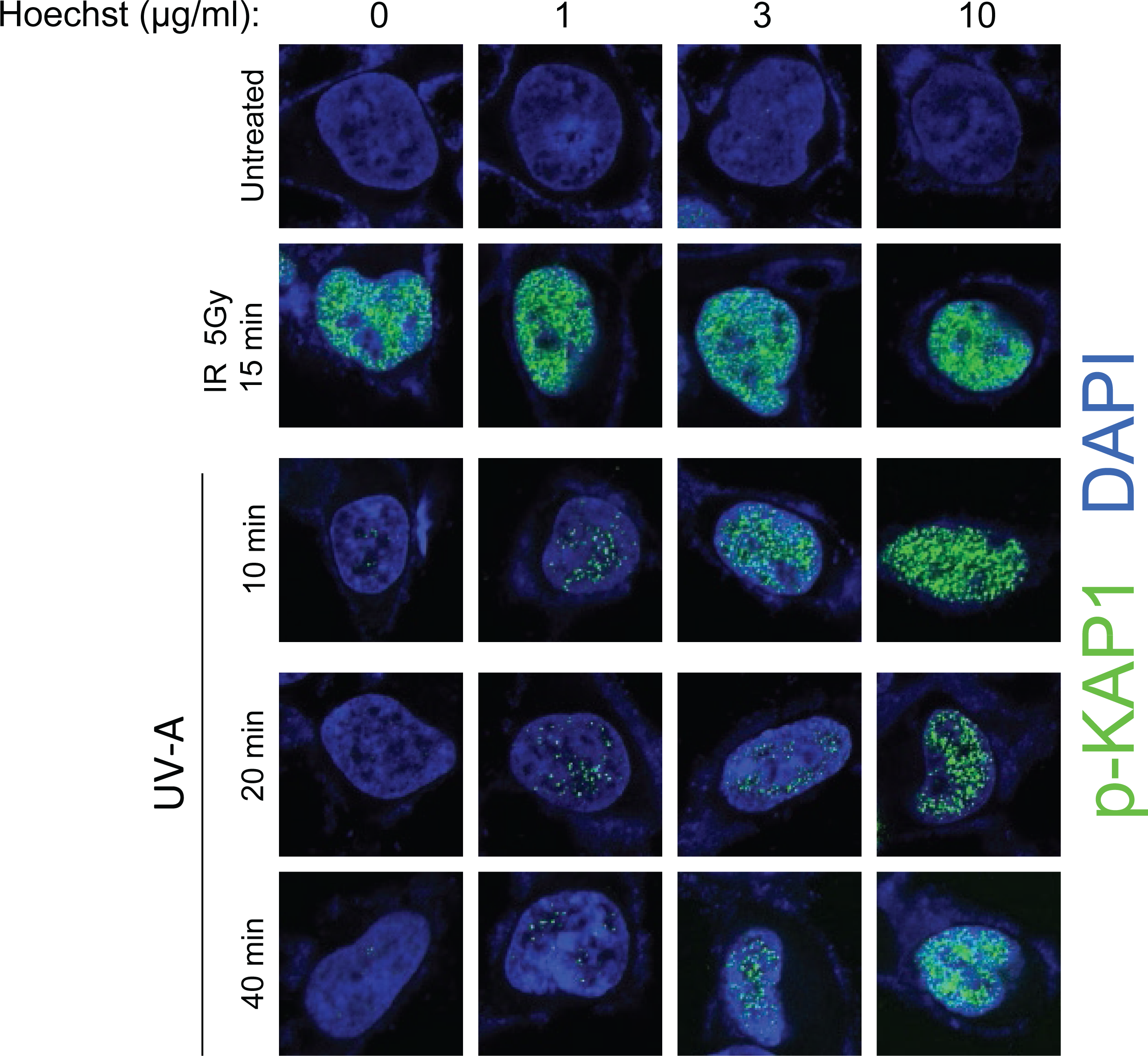
Hoechst 33258 pretreatment sensitizes cells to DNA damage. HEK293 cells were exposed to IR (5 Gy, 15 minutes) with or without Hoechst 33258 (1, 3, or 10 μg/ml; 3 hours) treatment or treated with UV-A (0.20 J/m^2^/s for 10, 20 or 40 minutes) with or without Hoechst 33258 (1, 3 or 10 μg/ml; 3 hours) treatment or treated with Hoechst 33258 alone (1, 3 or 10 μg/ml; 3 hours). Cells were immunostained with anti-pKAP1 (S824) and counterstained with DAPI. Images were acquired at 63x power.

**Supplemental Figure 9.**
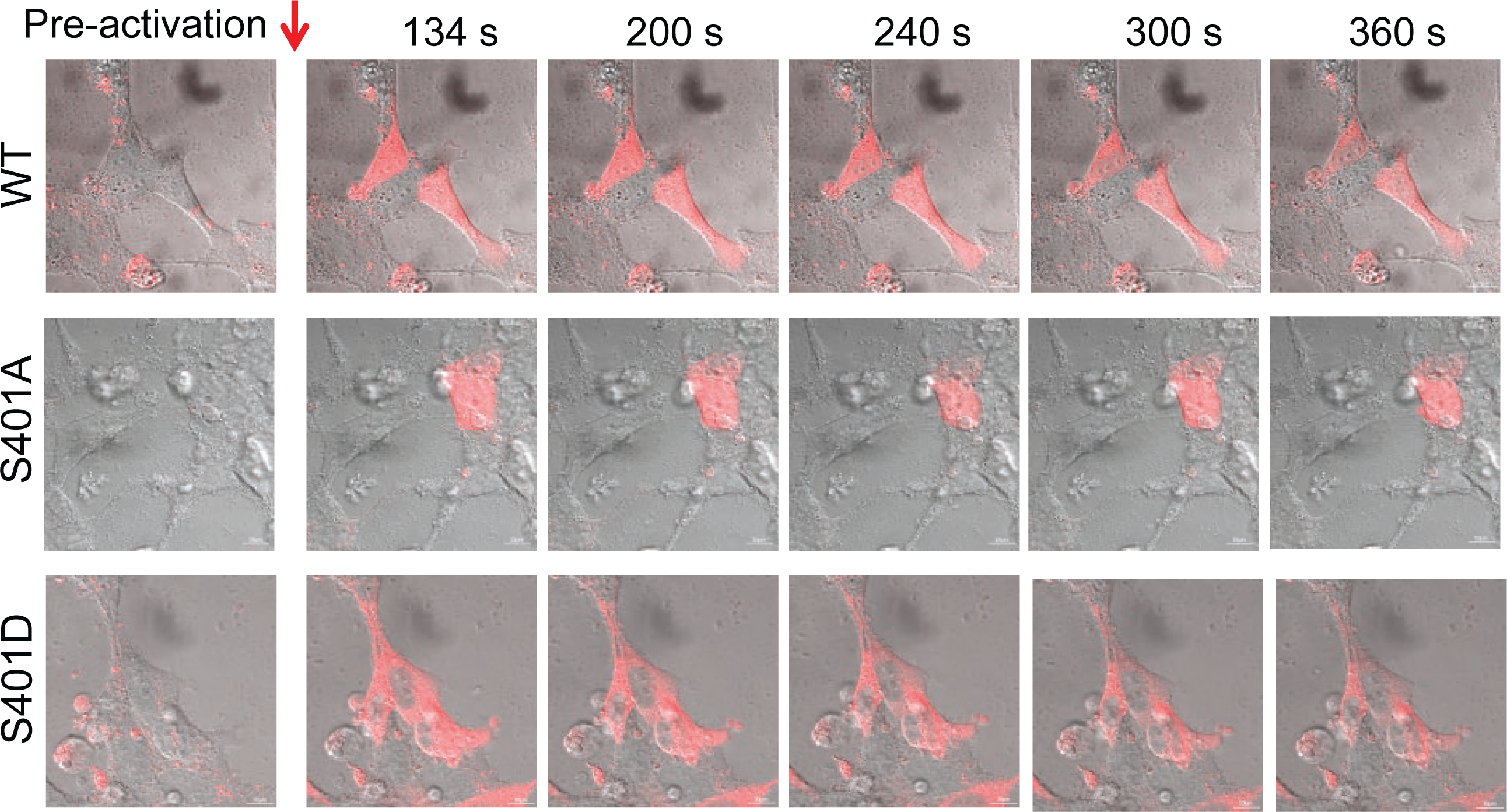
PR65 nuclear - cytoplasmic shuttling. Nuclear and cytoplasmic fluorescence over time. HEK293 cells were transfected with pPAmCherry-PR65 (WT, S401A and S401D) in a glass bottom dish. Forty-eight hours post-transfection sub-nuclear ROI (4 μm x 4 μm) were photo-activated by 405 nm laser and cells monitored over time by live cell imaging performed on Zeiss LSM 710 and images were acquired at 63x.

**Supplemental Figure 10.**
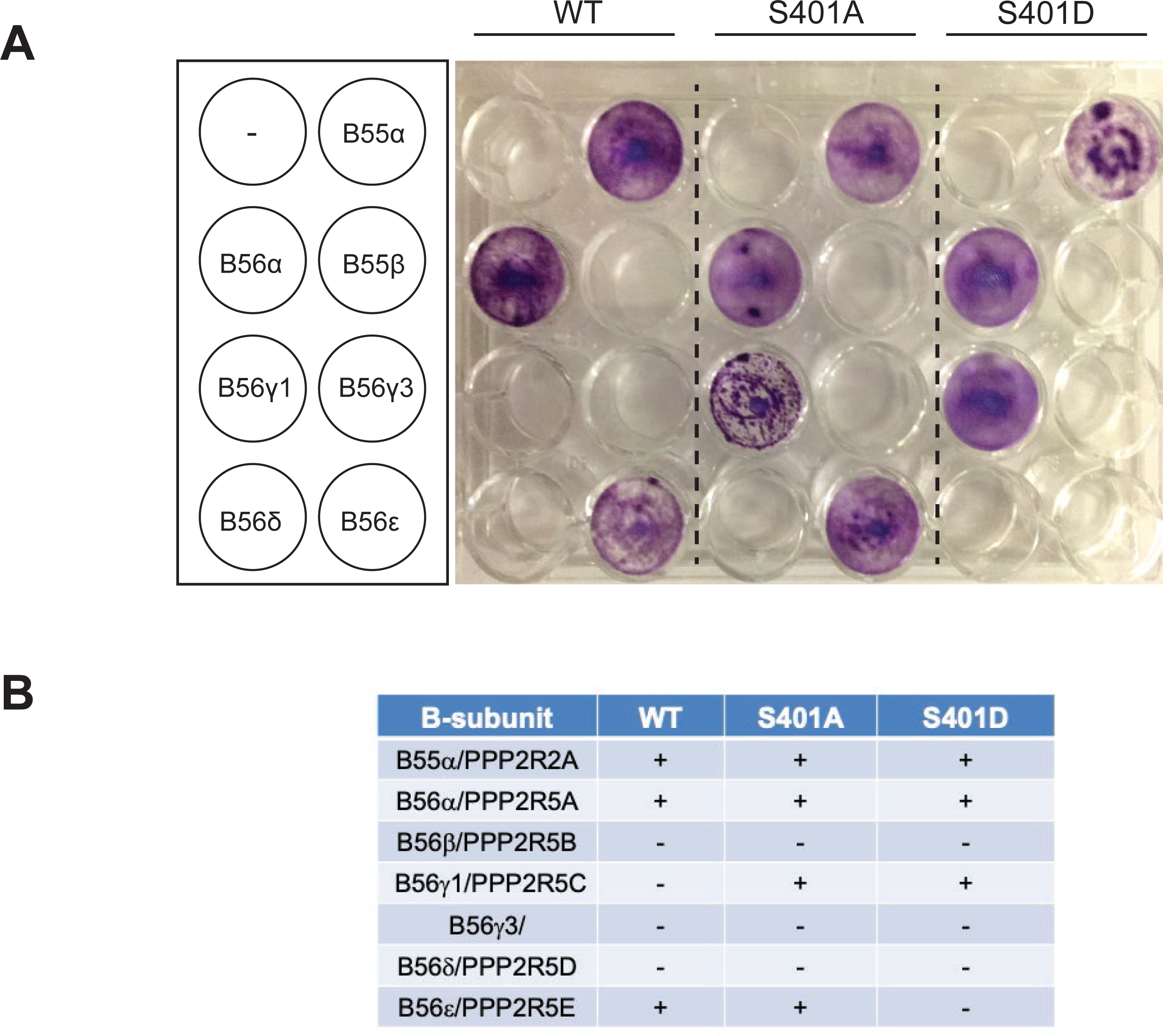
Tolerability of B subunit overexpression in PR65 MEFs. PR65-WT, -S401A, and -S401D MEFs were transfected with pCEP-4HA-B56 alpha, - B56 gamma3, -B56 gamma1, -B56 epsilon, -B56 delta, -B56 beta, or -B55 beta and selected for hygromycin (200 μg/ml) resistance. pCEP plasmids are maintained episomally as EBVori-EBNA plasmids in transfected MEFs (2).

**Supplemental Table 1.** Statistics for AFM studies. Four samples were amplified: wild-type (WT, *n* = 11,303 measured strands), S401A (*n* = 6,530), S401D (*n* = 12,160), and a synthetic fragment (Syn, *n* = 6,035) of equivalent length and sequence as WT. Gaussian distribution of 3 populations for WT, S01D and Syn and five populations for S401A was analyzed. Analysis of traced length distributions was performed by fitting Gaussian mixture models then analyzing the Bayesian information criteria to determine the number of Gaussian distributions that produced the best fit. The 95% confidence interval for the mean of each fitted Gaussian was determined using a bootstrapping approach with *n =* 10,000.

| Row | gaussian1 | gaussian2 | gaussian3 |
| --- | --- | --- | --- |
| mu | 112.6713334 | 47.99697261 | 172.1115854 |
| mu_CI_lower | 112.1608046 | 47.03594875 | 159.9828084 |
| mu_CI_upper | 113.1693958 | 48.97239987 | 186.0252576 |
| sigma | 20.57601541 | 13.99117091 | 94.11453542 |
| sigma_CI_lower | 20.13991044 | 13.10672902 | 85.39626477 |
| sigma_CI_upper | 21.02688002 | 14.89745807 | 102.5372019 |
| proportion | 0.8072826 | 0.151702945 | 0.041014455 |
| proportion_CI_lower | 0.796783762 | 0.142667002 | 0.034361264 |
| proportion_CI_upper | 0.817558008 | 0.16064956 | 0.047994046 |

S401A
| Row | gaussian1 | gaussian2 | gaussian3 | gaussian4 | gaussian5 |
| --- | --- | --- | --- | --- | --- |
| mu | 107.8401084 | 61.9504846 | 262.3366658 | 142.5177 | 48.63107 |
| mu_CI_lower | 106.6717451 | 60.16653504 | 203.624445 | 131.1313 | 44.22629 |
| mu_CI_upper | 115.4548081 | 68.23947331 | 377.3792976 | 165.2139 | 107.7963 |
| sigma | 21.32404405 | 9.207093964 | 124.878936 | 42.67979 | 21.57652 |
| sigma_CI_lower | 18.25635918 | 7.568911911 | 80.64580349 | 32.13473 | 8.859971 |
| sigma_CI_upper | 23.07417712 | 24.27469864 | 170.3866235 | 51.42518 | 43.28659 |
| proportion | 0.569086021 | 0.115927342 | 0.007074914 | 0.198092 | 0.109819 |
| proportion_CI_lower | 0.357837158 | 0.080946403 | 0.002256971 | 0.10822 | 0.042099 |
| proportion_CI_upper | 0.624558375 | 0.346681673 | 0.017940919 | 0.271828 | 0.196641 |

S401D
| Row | gaussian1 | gaussian2 | gaussian3 |
| --- | --- | --- | --- |
| mu | 106.0315687 | 46.76675558 | 174.8895841 |
| mu_CI_lower | 105.4597517 | 46.39652219 | 165.5553866 |
| mu_CI_upper | 106.6041131 | 47.13127314 | 185.3803763 |
| sigma | 20.54104028 | 9.570053569 | 93.10054925 |
| sigma_CI_lower | 19.98778326 | 9.266241556 | 87.48585699 |
| sigma_CI_upper | 21.09092988 | 9.868076039 | 98.51755827 |
| proportion | 0.607900591 | 0.335846752 | 0.056252658 |
| proportion_CI_lower | 0.597047248 | 0.326093943 | 0.049496932 |
| proportion_CI_upper | 0.619078528 | 0.345248898 | 0.063379692 |

Syn
| Row | gaussian1 | gaussian2 | gaussian3 |
| --- | --- | --- | --- |
| mu | 119.0488805 | 44.52886136 | 171.5183648 |
| mu_CI_lower | 118.570992 | 41.67076681 | 161.6553723 |
| mu_CI_upper | 119.597509 | 47.40926267 | 235.400503 |
| sigma | 18.32874438 | 20.95980558 | 91.48017913 |
| sigma_CI_lower | 17.97499435 | 18.70377407 | 27.44910405 |
| sigma_CI_upper | 18.77843553 | 23.45894998 | 129.842548 |
| proportion | 0.925191784 | 0.055257503 | 0.019550713 |
| proportion_CI_lower | 0.91788522 | 0.049143018 | 0.007864581 |
| proportion_CI_upper | 0.937139109 | 0.064220123 | 0.02474155 |

**Supplemental Table 2.**
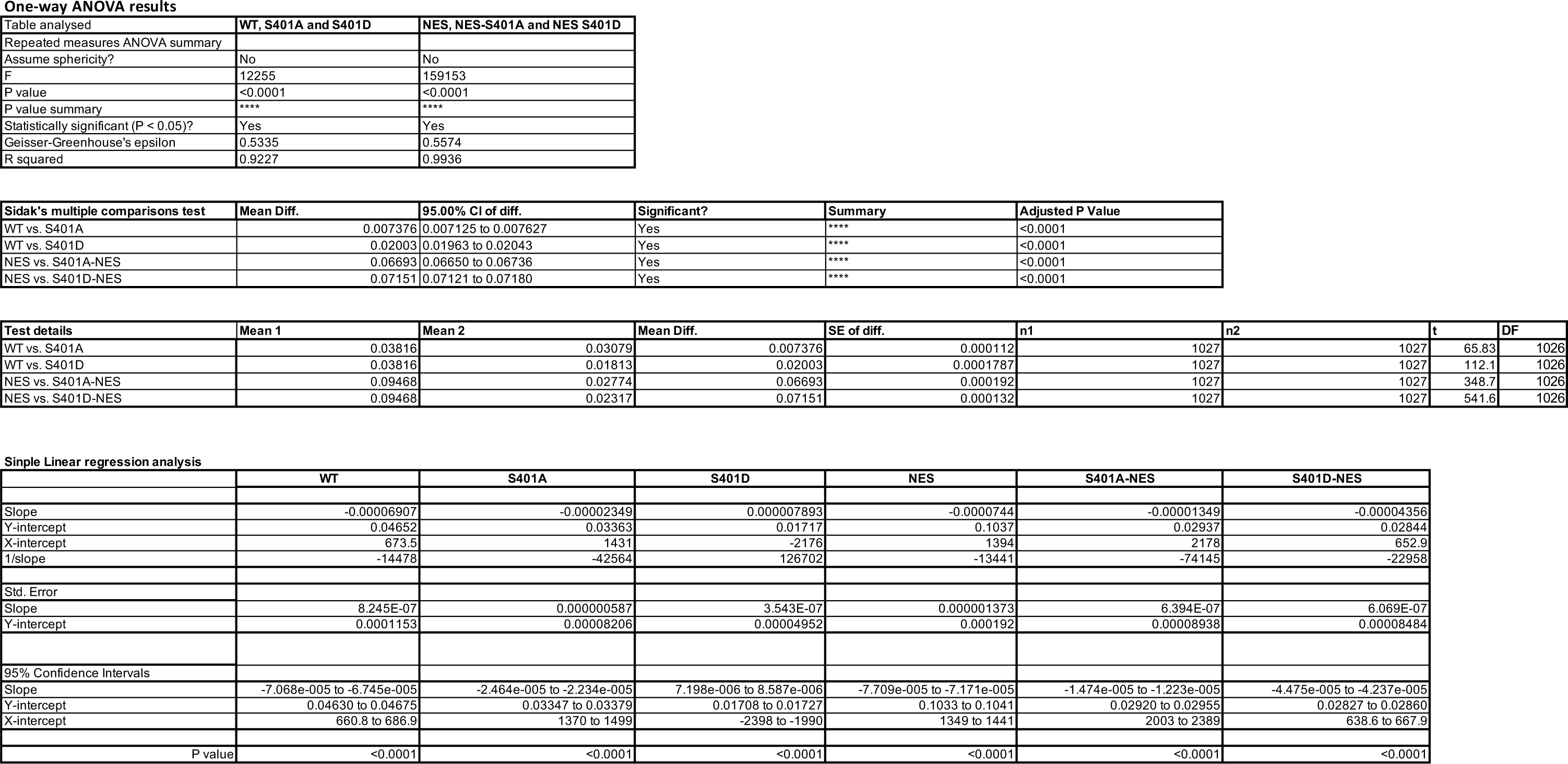
Statistics for photoactivation experiment (Figure 9E). One way ANOVA test was conducted for the nuclear to cytoplasmic intensity ratios in the time series studies for WT, S401A and S40D as well as NES, NES-S401A and NES-S401D groups. On Multiple comparison test, the differences between WT vs S401A, WT vs S401D, NES vs NES-S401A and NES vs NEs-S401D were highly significant. We also performed simple linear regression on the time series of the N/C intensities and found that the slopes of WT, S401A and S401d as well as NES, NES-S401A and NES - S401D were significant different than each other.

|  |  |  |
| --- | --- | --- |
| Table analysed | WT, S401A and S401D | NES, NES-S401A and NES S401D |
| Repeated measures ANOVA summary |  |  |
| Assume sphericity? | No | No |
| F | 12255 | 159153 |
| P value | <0.0001 | <0.0001 |
| P value summary | **** | **** |
| Statistically significant (P < 0.05)? | Yes | Yes |
| Geisser-Greenhouse's epsilon | 0.5335 | 0.5574 |
| R squared | 0.9227 | 0.9936 |

|  |  |  |  |  |  |
| --- | --- | --- | --- | --- | --- |
| Sidak's multiple comparisons test | Mean Diff. | 95.00% CI of diff. | Significant? | Summary | Adjusted P Value |
| WT vs. S401A | 0.007376 | 0.007125 to 0.007627 | Yes | **** | <0.0001 |
| WT vs. S401D | 0.02003 | 0.01963 to 0.02043 | Yes | **** | <0.0001 |
| NES vs. S401A-NES | 0.06693 | 0.06650 to 0.06736 | Yes | **** | <0.0001 |
| NES vs. S401D-NES | 0.07151 | 0.07121 to 0.07180 | Yes | **** | <0.0001 |

|  |  |  |  |  |  |  |  |  |
| --- | --- | --- | --- | --- | --- | --- | --- | --- |
| Test details | Mean 1 | Mean 2 | Mean Diff. | SE of diff. | n1 | n2 | t | DF |
| WT vs. S401A | 0.03816 | 0.03079 | 0.007376 | 0.000112 | 1027 | 1027 | 65.83 | 1026 |
| WT vs. S401D | 0.03816 | 0.01813 | 0.02003 | 0.0001787 | 1027 | 1027 | 112.1 | 1026 |
| NES vs. S401A-NES | 0.09468 | 0.02774 | 0.06693 | 0.000192 | 1027 | 1027 | 348.7 | 1026 |
| NES vs. S401D-NES | 0.09468 | 0.02317 | 0.07151 | 0.000132 | 1027 | 1027 | 541.6 | 1026 |

Simple Linear regression analysis
|  |  |  |  |  |  |  |
| --- | --- | --- | --- | --- | --- | --- |
|  | WT | S401A | S401D | NES | S401A-NES | S401D-NES |
| Slope | -0.00006907 | -0.00002349 | 0.000007893 | -0.0000744 | -0.00001349 | -0.00004356 |
| Y-intercept | 0.04652 | 0.03363 | 0.01717 | 0.1037 | 0.02937 | 0.02844 |
| X-intercept | 673.5 | 1431 | -2176 | 1394 | 2178 | 652.9 |
| 1/slope | -14478 | -42564 | 126702 | -13441 | -74145 | -22958 |
| Std. Error |  |  |  |  |  |  |
| Slope | 8.245E-07 | 0.000000587 | 3.543E-07 | 0.000001373 | 6.394E-07 | 6.069E-07 |
| Y-intercept | 0.0001153 | 0.00008206 | 0.00004952 | 0.000192 | 0.00008938 | 0.00008484 |
| 95% Confidence Intervals |  |  |  |  |  |  |
| Slope | -7.068e-005 to -6.745e-005 | -2.464e-005 to -2.234e-005 | 7.198e-006 to 8.587e-006 | -7.709e-005 to -7.171e-005 | -1.474e-005 to -1.223e-005 | -4.475e-005 to -4.237e-005 |
| Y-intercept | 0.04630 to 0.04675 | 0.03347 to 0.03379 | 0.01708 to 0.01727 | 0.1033 to 0.1041 | 0.02920 to 0.02955 | 0.02827 to 0.02860 |
| X-intercept | 660.8 to 686.9 | 1370 to 1499 | -2398 to -1990 | 1349 to 1441 | 2003 to 2389 | 638.6 to 667.9 |
| P value | <0.0001 | <0.0001 | <0.0001 | <0.0001 | <0.0001 | <0.0001 |

